# *In Vitro* Modeling of CD8 T Cell Exhaustion Enables CRISPR Screening to Reveal a Role for BHLHE40

**DOI:** 10.1101/2023.04.17.537229

**Authors:** Jennifer E. Wu, Sasikanth Manne, Shin Foong Ngiow, Amy E. Baxter, Hua Huang, Elizabeth Freilich, Megan L. Clark, Joanna H. Lee, Zeyu Chen, Omar Khan, Ryan P. Staupe, Yinghui J. Huang, Junwei Shi, Josephine R. Giles, E. John Wherry

## Abstract

Identifying novel molecular mechanisms of exhausted CD8 T cells (T_ex_) is a key goal of improving immunotherapy of cancer and other diseases. However, high-throughput interrogation of *in vivo* T_ex_ can be costly and inefficient. *In vitro* models of T_ex_ are easily customizable and quickly generate high cellular yield, offering an opportunity to perform CRISPR screening and other high-throughput assays. We established an *in vitro* model of chronic stimulation and benchmarked key phenotypic, functional, transcriptional, and epigenetic features against bona fide *in vivo* T_ex_. We leveraged this model of *in vitro* chronic stimulation in combination with pooled CRISPR screening to uncover transcriptional regulators of T cell exhaustion. This approach identified several transcription factors, including BHLHE40. *In vitro* and *in vivo* validation defined a role for BHLHE40 in regulating a key differentiation checkpoint between progenitor and intermediate subsets of T_ex_. By developing and benchmarking an *in vitro* model of T_ex_, we demonstrate the utility of mechanistically annotated *in vitro* models of T_ex_, in combination with high-throughput approaches, as a discovery pipeline to uncover novel T_ex_ biology.

## INTRODUCTION

T cell exhaustion is a distinct cell state that arises during chronic viral infection and cancer. Exhausted CD8 T cells (T_ex_) are defined by reduced proliferative potential, high and sustained expression of inhibitory receptors (IRs), decreased production of effector cytokines, and a distinct transcriptional and epigenetic program (*1*). The inability to durably reverse exhaustion represents a major barrier to treatment of cancers and chronic viral infections. Efforts to develop more effective therapies for these diseases are inhibited by a limited understanding of the molecular mechanisms that underlie T_ex_ biology. Despite a substantial body of work analyzing T_ex_ biology in search of novel therapeutic targets, most of these efforts have been relatively low-throughput due to limitations of *in vivo* mouse models. Defining scenarios in which key aspects of T_ex_ biology can be modulated and even screened in a high-throughput fashion could reveal new therapeutic opportunities.

Mouse models of chronic viral infection, such as lymphocytic choriomeningitis virus (LCMV), and cancer have been instrumental in establishing our understanding of CD8 T cell exhaustion, including discoveries with therapeutic relevance such as checkpoint pathway blockade (*2–4*). Despite their utility, *in vivo* models have limitations in efficiency and cellular yield. Mouse models of T cell exhaustion often generate relatively small numbers of T_ex_, limiting approaches that require large cell numbers. As a result, high-throughput experimental assays are challenging due to differences in scale between cells required and cells available. For example, shRNA or CRISPR screens in *in vivo* models require pooling of large numbers of mice to obtain sufficiently high cell/guide numbers to achieve robust screening results. Increasing the input cell number in the interest of maximizing yield can perturb pathogenesis and alter cellular differentiation. Adoptive transfer of a higher number of LCMV-specific CD8 T cells results in faster control of viral replication and/or increased immunopathology (*5–7*) and can influence CD8 T cell differentiation (*8–10*), circumventing the very biology of T cell exhaustion that is the intent of the study. Thus, although *in vivo* CRISPR screening models have uncovered valuable new insights into T_ex_ and other T cell states, overcoming limitations in scale would allow additional layers of biology to be interrogated.

*In vitro* models are an attractive alternative to *in vivo* mouse models for screening and other discovery platforms because cellular output can be generated quickly and efficiently. *In vitro* models can be scaled up to maximize cellular yield and/or large amounts of cellular material for high-throughput assays (e.g. proteomics, metabolomics). However, modeling complex *in vivo* phenomena *in vitro* also has inherent challenges. *In vitro* models cannot fully recapitulate the complexities and nuance of *in vivo* T_ex_ biology: for example, 3D tissue architecture and cell homing/migration, are challenging to model *in vitro*. Yet the reductionist nature of *in vitro* models provides a unique opportunity to dissect specific pathways. *In vitro* cultures can be easily customized to interrogate the causal effects of a diverse range of stimuli or conditions, such as cytokines and other secreted molecules, hypoxia, metabolites, pharmacological small molecules, or even genetic manipulations. Regardless, the utility of any model depends on an accurate evaluation of which features of *in vivo* biology are effectively modeled *in vitro* and which are not. Detailed benchmarking and analysis of *in vitro* models of T_ex_ could have considerable value given the importance of this cell type in human disease.

Sustained TCR engagement is a central driver of CD8 T cell exhaustion *in vivo* (*11–13*). Repeated or continuous TCR stimulation *in vitro* can also successfully induce phenotypic features of T_ex_, such as loss of proliferative potential (*14*), expression of PD-1 and other IRs (*15*), reduced effector function (*16*), or a combination of these properties (*17–21*). Collectively, these studies demonstrate the utility of *in vitro* models approximating T cell exhaustion. However, defining how much of the overall program of *in vivo* T_ex_ biology is accurately captured in these settings remains a challenge. A more detailed understanding of which aspects of T_ex_ biology are and are not capable of being modeled *in vitro* will enable more successful downstream application of *in vitro* models and screens to identify underlying mechanisms and therapeutic opportunities.

To address these issues, we developed an *in vitro* model that approximates T_ex_ via chronic administration of cognate peptide in a manner intended to achieve continuous antigenic stimulation. We benchmarked these *in vitro* chronically stimulated CD8 T cells against *in vivo* T_ex_ generated during chronic LCMV infection to confirm that phenotypic, functional, transcriptional, and epigenetic features of T_ex_ were appropriately recapitulated *in vitro*. We then leveraged this *in vitro* culture model in combination with CRISPR/Cas9 screening of transcriptional pathways to identify a role for the transcription factor (TF) BHLHE40 in T cell exhaustion. We returned to mouse models of chronic viral infection to interrogate the function of BHLHE40 in *in vivo* T_ex_ and found that BHLHE40 regulates a differentiation checkpoint between progenitor T_ex_ and intermediate and terminally differentiated subsets of T_ex_. Our data suggest that biology targeted by BHLHE40 may be relevant for therapeutically manipulating these T_ex_ subsets involved in response to checkpoint blockade.

Here we demonstrate the utility of mechanistically annotated *in vitro* models of T cell exhaustion. In combination with CRISPR technology, this *in vitro* model of chronic stimulation enables high-throughput screening of transcriptional regulators of T_ex_, after which individual hits can be further interrogated *in vitro* and *in vivo*. This foundation of *in vitro* modeling of T_ex_ will allow scaling of other discovery approaches to probe deeper into the mechanisms of CD8 T cell exhaustion.

## RESULTS

### Chronic antigenic stimulation *in vitro* induces the molecular phenotype of CD8 T cell exhaustion

Many *in vitro* models recapitulate some of the known features of T_ex_ (*14, 17-19, 21*). However, it is often unclear which features or biological modules of *in vivo* biology can and cannot be recreated *in vitro*. To address this question, we aimed first to develop an *in vitro* model of T_ex_ and subsequently to deeply interrogate and benchmark the phenotypic, functional, transcriptional, and epigenetic features of this model in comparison to *in vivo* T_ex_ generated during chronic LCMV infection.

We established an *in vitro* model of T_ex_, building on previous work from Beltra et al (*15*). Naïve CD8 T cells were isolated from mice transgenic for a TCR that recognizes the D^b^GP^33-41^ epitope of LCMV (P14 mice). These P14 cells were co-cultured with dendritic cells from WT C57BL/6 mice that were pulsed with D^b^GP^33-41^ peptide. To mimic chronic antigenic stimulation characteristic of that which drives T_ex_ differentiation *in vivo*, P14 cells were further stimulated with D^b^GP^33-41^ peptide and IL-2 every 2 days thereafter (Fig. 1A). This timing was chosen based on previous work showing that peptide stimulation under *in vitro* conditions lasts ∼2 days or less (*22*). We also generated an *in vitro* acutely stimulated condition, in which P14 cells received D^b^GP^33-41^ peptide only at day 0. At day 7 of *in vitro* culture, P14 cells were harvested and analyzed for activation and differentiation by flow cytometry. As a benchmark for bona fide T_ex_ biology, *in vivo* CD8 T_ex_ were generated via LCMV clone 13 (Cl13) infection and analyzed in parallel.

**Fig. 1.**
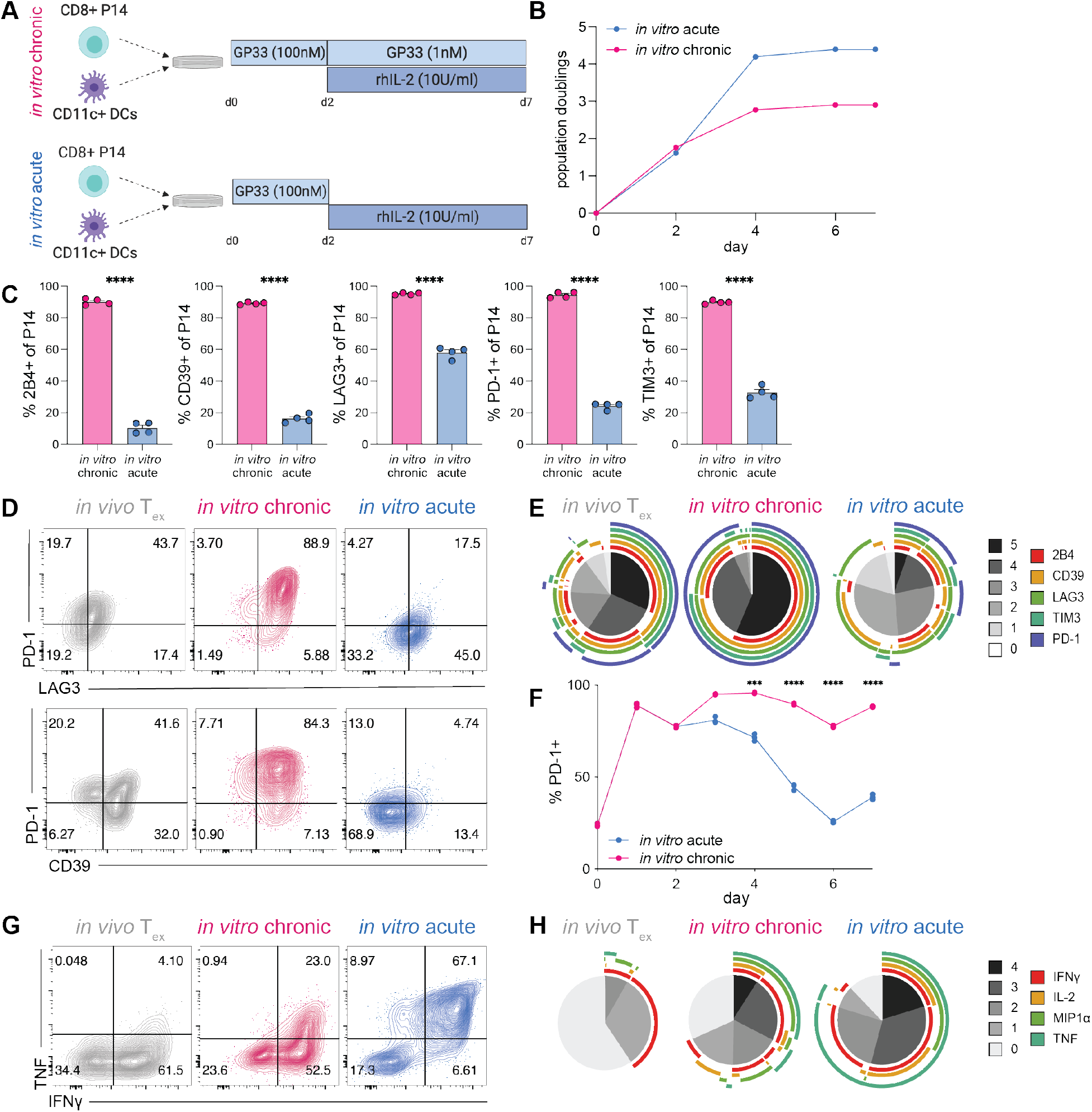
Chronic antigenic stimulation *in vitro* induces key features of T_ex_. **(A)** Experiment schematic of chronic and acute stimulation of P14 cells *in vitro*. **(B)** Cell expansion during chronic and acute stimulation *in vitro*. **(C)** Percent expression of IRs by chronically and acutely stimulated P14 cells (gated on CD44^hi^ CD8+ live singlets); two technical replicates of two biological replicates shown. Significance calculated by unpaired two-tailed t test; **\*\*\*\***p<0.0001. **(D)** Representative flow cytometry data and **(E)** SPICE analysis of IR co-expression by *in vivo* T_ex_ (LCMV-Cl13 30dpi), *in vitro* chronically stimulated P14 cells, and *in vitro* acutely stimulated P14 cells (gated on CD44^hi^ CD8+ live singlets). **(F)** Longitudinal PD-1 expression on *in vitro* chronically and acutely stimulated P14 cells. Representative of 2 experiments; significance calculated by paired two-tailed t test; **\***p<0.05, **\*\***p<0.01, **\*\*\***p<0.001, **\*\*\*\***p<0.0001. **(G)** Representative flow cytometry data and **(H)** SPICE analysis of co-production of effector cytokines in *in vivo* T_ex_ (LCMV-Cl13 30dpi), *in vitro* chronically stimulated P14 cells, and *in vitro* acutely stimulated P14 cells. **(B-E, G-H)** Representative of >3 experiments. **(D,G)** Numbers in flow cytometry plots indicate percentage of parent population within each gate. **(E,H)** Grayscale sections in SPICE plots indicate number of IRs/cytokines co-expressed; colored bands indicate individual IRs/cytokines.

*In vivo* T_ex_ are phenotypically distinct from naïve (T_n_), effector (T_eff_) and memory (T_mem_) CD8 T cells and are characterized by: 1) decreased proliferative potential, 2) high and stable expression of inhibitory receptors (IRs), and 3) decreased production of effector cytokines (*11, 12, 23*). We benchmarked our *in vitro* model against these known hallmarks of *in vivo* T_ex_, beginning with proliferation and expansion. Both *in vitro* conditions induced proliferation and expansion between days 0 and 4 of culture, after which proliferation slowed and eventually plateaued. However, whereas acutely stimulated P14 cells reached 4 population doublings, chronically stimulated P14 cells were unable to maintain similar levels of expansion and plateaued at less than 3 population doublings, despite continuous exposure to antigenic stimulation (Fig. 1B). These data suggest that persistent antigenic stimulation is sufficient to reduce proliferative expansion and/or potential. To interrogate the contribution of proliferation and cell death to this overall decreased expansion, we assessed expression of the cell cycle protein KI67 and pro- and anti-apoptotic proteins BIM and BCL-2, respectively. KI67 expression was maintained between days 4-7 of *in vitro* culture; however, this ongoing cell cycle was accompanied by an increasing BIM/BCL-2 ratio, indicating that chronic stimulation results in sustained proliferation but also increased sensitivity to cell death, leading to high cell turnover and a net overall lack of expansion in cell numbers (Fig. S1A). These results are consistent with observations *in vivo* T_ex_, for which ongoing stimulation is associated with continued cell cycle and proliferation, but no net increase in cell numbers (*24–31*).

Chronic antigen stimulation *in vitro* was sufficient to induce high expression of multiple IRs, characteristic of *in vivo* T_ex_ (*12, 32*). Whereas acutely stimulated P14 cells demonstrated significantly lower expression of IRs, such as PD-1 and LAG3, than *in vivo* T_ex_, chronic stimulation induced high expression of IRs as a percentage (Fig. 1C) and on a per cell level, as quantified by MFI (Fig. S1B). Furthermore, a higher proportion of chronically stimulated P14 cells co-expressed combinations of IRs, including PD-1 and LAG3 or CD39 (Fig. 1D), than acutely stimulated P14 cells or even *in vivo* T_ex_. Because increased diversity and co-expression of IRs is associated with progression toward terminal exhaustion (*26, 32, 33*), we performed SPICE analysis to assess IR co-expression. More than 50% of chronically stimulated P14 cells co-expressed all 5 IRs analyzed, an even higher proportion of IR co-expression than observed on *in vivo* T_ex_ (∼30%) (Fig. 1E). In contrast, less than 10% of acutely stimulated P14 cells co-expressed 5 or more IRs. Furthermore, whereas acute stimulation induced IR expression shortly after activation that waned over time, high expression of IRs was maintained on P14 cells that had undergone chronic stimulation *in vitro* (Fig. 1F), consistent with stable maintenance of IR expression by *in vivo* T_ex_. We compared this antigen-specific *in vitro* model of chronic stimulation to a previously published model of chronic stimulation using non-antigen-specific TCR engagement (*18*). Although IR expression was similar between both *in vitro* models, chronic stimulation via αCD3/αCD28 resulted in substantially decreased expansion and cellular yield (Fig. S1C). Also, to assess whether additional rounds of antigen administration would further enhance features of T_ex_, we extended our *in vitro* chronic stimulation by an additional 3 days. Chronically stimulated P14 cells at d10 of *in vitro* culture had comparable IR expression to those at day 7, suggesting little increase in exhaustion phenotype; however, extending chronic stimulation *in vitro* to 10 days resulted in poorer cell recovery (Fig. S1D).

We next evaluated our *in vitro* generated P14 populations for reduced and/or altered effector cytokine production characteristic of T_ex_ (*11, 12, 23*). Whereas *in vitro* acutely stimulated P14 cells produced multiple effector cytokines in response to peptide restimulation, chronically stimulated P14 cells had a reduced capacity to produce TNF and IFNγ, though the extent of loss of cytokine production was not as severe as that of *in vivo* T_ex_ (Fig. 1G). Because hierarchical loss of effector cytokine production is associated with increased severity of exhaustion (*11, 12*), we assessed cytokine co-production via SPICE analysis. Approximately 20% of P14 cells were able to co-produce all 4 effector cytokines analyzed after acute stimulation *in vitro*. This multi-functional population was substantially reduced in *in vitro* chronically stimulated P14 cells to approximately 10% and nearly absent in *in vivo* T_ex_ (Fig. 1H). Collectively, these data suggest that chronic stimulation *in vitro* was sufficient to recapitulate known cellular hallmarks of T_ex_ biology, including reduced proliferative potential, sustained high expression and co-expression of multiple IRs, and decreased production of effector cytokines.

### *In vitro* chronically stimulated CD8 T cells develop a transcriptional signature of T_ex_

In addition to phenotypic and functional properties, *in vivo* generated T_ex_ are defined by unique transcriptional and epigenetic landscapes distinct from those of T_n_, T_eff_, and T_mem_ (*1, 12, 34-38*). To assess global transcriptional changes induced by acute versus chronic stimulation *in vitro*, we analyzed P14 cells by RNA-seq on days 0, 4, and 7 of *in vitro* culture. By number of differentially expressed genes (DEGs), the most substantial differences were observed between naïve (d0) and *in vitro* stimulated P14 cells at d4 regardless of stimulation condition (Fig. 2A), indicating that the most robust transcriptional changes occurred upon the transition from naïve to activated CD8 T cell. Whereas acute and chronically stimulated P14 cells only differed by 634 DEGs at d4 of *in vitro* culture, this number grew to 2844 by d7, suggesting that acute and chronic stimulation induced similar early transcriptional programs that diverged significantly by d7 (Fig. 2A). Furthermore, unbiased DEG analysis identified a cluster of genes uniquely expressed at d7 of chronic stimulation, including *Havcr2*, *Ccl3*, and *Xcl1* (Fig. 2B). Consistent with these observations, principal component analysis (PCA) revealed that acutely and chronically stimulated P14 cells clustered together at d4 (Fig. S2A). By d7, however, acutely and chronically stimulated P14 cells occupied distinct principal component space from all other *in vitro* generated populations. Collectively, these data indicate that, despite some overlap in transcriptional programs shortly after activation, acute and chronic stimulation *in vitro* generate unique and divergent transcriptional states by d7. Furthermore, this transcriptional divergence between d4 and d7 of *in vitro* culture suggests coordinated temporal regulation of gene expression in response to both acute and chronic stimulation *in vitro*.

**Fig. 2.**
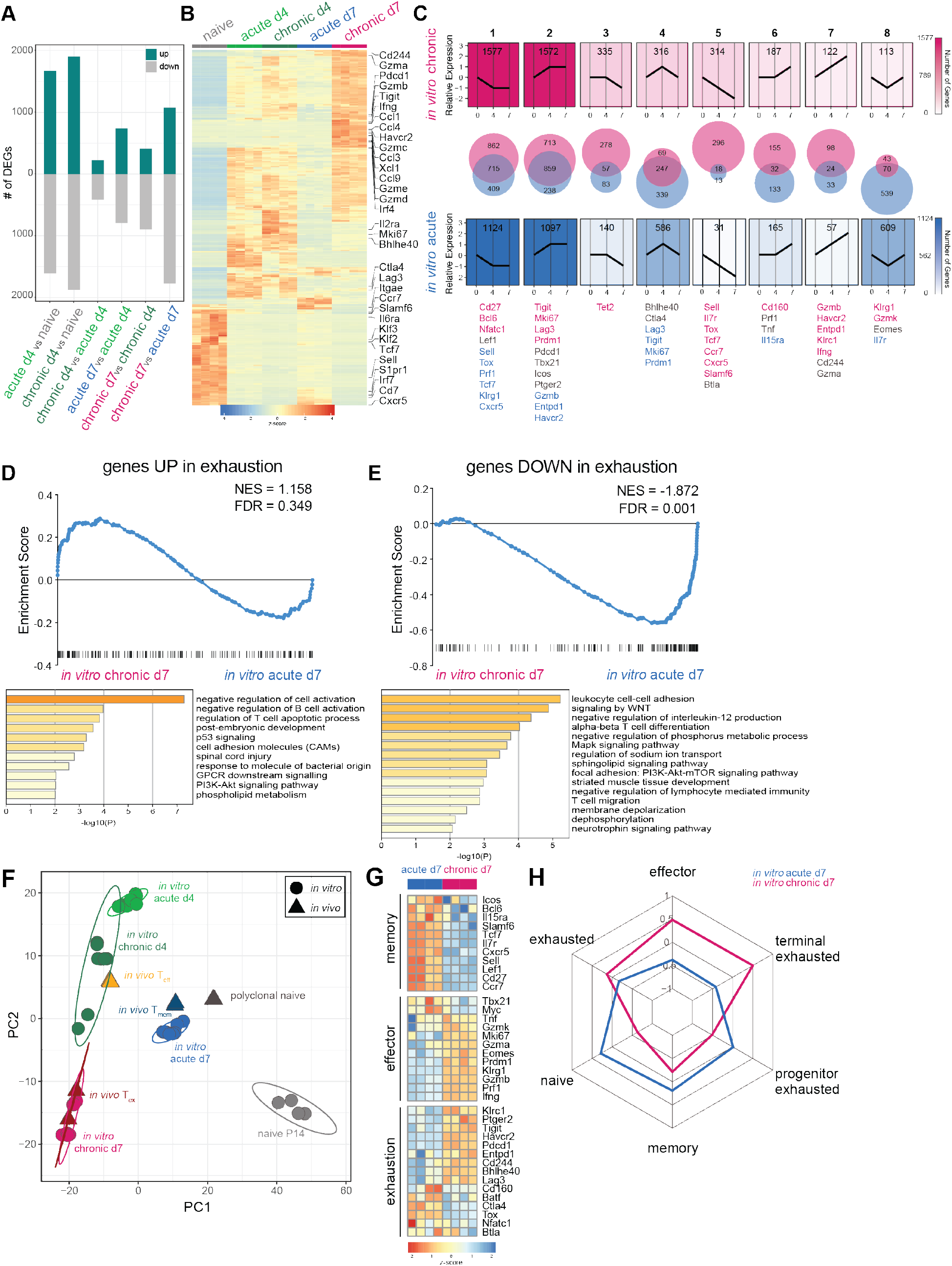
*In vitro* chronically stimulated P14 cells develop a transcriptional signature of T_ex_. **(A)** Number of differentially expressed genes (DEGs; filtered on log_2_ fold change (lfc)>1 and p<0.05) between pairwise comparisons as indicated. **(B)** Top DEGs (filtered on lfc>6; p<0.01) from all pairwise comparisons between *in vitro* generated CD8 T cell subsets. **(C)** Trajectory analysis (see Methods) of gene expression patterns during chronic or acute stimulation *in vitro*. **(D-E)** [top] Gene set enrichment analysis (GSEA) of genes **(D)** upregulated and **(E)** downregulated in *in vivo* T_ex_ (*41*) after chronic or acute stimulation *in vitro*. [bottom] Gene ontology (GO) analysis of leading edge genes. **(F)** Principal component analysis (PCA) of RNA-seq data from *in vitro* chronically and acutely stimulated P14 cells and previously published *in vivo* CD8 T cell subsets (*37*). **(G)** Gene expression of manually curated lists of genes associated with T_mem_, T_eff_, or T_ex_ in *in vitro* chronically and acutely stimulated P14 cells. **(H)** Gene set variation analysis (GSVA) indicating comparative enrichment for various gene signatures (*28, 41*) after chronic or acute stimulation *in vitro*.

To further interrogate temporal transcriptional trajectories induced by acute and chronic stimulation *in vitro*, we manually binned genes into patterns of relative expression at days 0, 4, and 7 of *in vitro* culture (Fig. 2C). The two most common temporal patterns of gene expression in both acutely and chronically stimulated P14 were a decrease or increase from d0 to d4 followed by stable maintenance of expression from d4 to d7 (patterns 1 and 2, respectively). Whereas some genes within these patterns biased either toward chronic or acute stimulation, most genes were shared between both conditions. These two most common temporal gene patterns confirm that the greatest magnitude of change in gene expression occur upon the transition from naïve to activated CD8 T cell and highlight common biology found in both *in vitro* conditions.

To investigate transcriptional trajectories preferentially associated with acute stimulation, we examined genes in patterns 4 (transient upregulation at d4) and 8 (transient downregulation at d4) (Fig. 2C). Pattern 4 contained 586 genes that transiently increased during acute stimulation. Of these, less than half overlapped between acute and chronic stimulation; 339 of these changes were found only after acute stimulation. Genes in pattern 4 preferentially responsive to acute stimulation included *Lag3*, *Tigit*, and *Mki67*, consistent with acutely stimulated P14 cells becoming transiently activated before returning to a relative state of quiescence. Conversely, gene expression of *Lag3*, *Tigit*, and *Mki67* remained elevated in chronically stimulated P14 cells at d7 (pattern 2), consistent with a continued role for these genes in T_ex_. Pattern 8, also biased toward acute stimulation conditions, displayed an inverse pattern, with genes decreasing in expression at d4 then returning to baseline by d7. Of these 600 transcriptional changes biased toward acute stimulation, less than 100 were shared with chronic stimulation. For example, expression of *Il7r*, which encodes the IL-7 receptor crucial for homeostatic proliferation in T_n_ and T_mem_, followed this pattern of transient decrease. Thus, acute stimulation *in vitro* recapitulated key temporal changes in gene expression reminiscent of *in vivo* T cell activation and transition to a post-effector state.

To interrogate transcriptional trajectories preferentially associated with chronic stimulation, we examined genes that decreased (pattern 5) or increased (pattern 7) in expression throughout the course of *in vitro* culture (Fig. 2C). Pattern 5 was dominated by 296 genes biased toward chronic stimulation, with only 18 genes shared with acute stimulation. Genes in this pattern included *Sell*, *Il7r*, *Tcf7*, *Cxcr5*, and *Slamf6*, genes typically associated with stemness and quiescence. Pattern 7 included only 24 genes shared by both conditions and 98 genes biased toward chronic stimulation, including genes encoding IRs, such as *Havcr2* (TIM3) and *Entpd1* (CD39), as well as genes related to effector function, such as *Ifng* and *Gzmb*. Increased transcription despite decreased protein expression of effector molecules is a pattern consistent with early transcriptional studies of T_ex_ that first noted this disconnect between effector function and transcription of effector genes (*12*). Collectively, these patterns suggest differential regulation of transcriptional trajectories in response to chronic or acute stimulation *in vitro*. Furthermore, genes within temporal patterns preferentially responsive to chronic stimulation *in vitro* are associated with transcriptional evidence of persistent activation as well as robust downregulation of genes associated with durability, stemness, and other programs associated with T_mem_ development.

To quantify how gene expression induced by chronic stimulation *in vitro* compared to that of *in vivo* T_ex_, we employed gene set enrichment analysis (GSEA) (*39, 40*) of a previously published transcriptional signature of exhaustion (*41*). Genes upregulated in *in vivo* T_ex_ trended toward enrichment in *in vitro* chronically stimulated P14 cells (Fig. 2D), although this enrichment did not reach statistical significance. One possible interpretation of this result is that chronic stimulation *in vitro* recapitulates some, but not all, features of *in vivo* T_ex_. It is also possible that a strong transcriptional signature of activation in this *in vitro* model obscures significant enrichment of this exhaustion signature. To investigate which transcriptional pathways from this *in vivo* T_ex_ gene signature were and were not captured by chronic stimulation *in vitro*, we focused on the leading edge of enriched genes and applied gene ontology (GO) analysis. Leading edge genes included known exhaustion-related genes, such as *Pdcd1*, *Lag3*, *Tigit,* and *Prdm1* (Fig. S2B). GO analysis of these leading edge genes highlighted multiple pathways involved in negative regulation of T cell activation (Fig. 2D). These data were supported by the observation that a gene signature of *in vitro* activation was more highly enriched in chronically stimulated cells at d4 compared to d7 (Fig. S2C). Conversely, the *in vivo* T_ex_ gene signature enriched in chronically stimulated cells at d7 compared to d4 (Fig. S2D). Collectively, these GSEA analyses indicate an overall transcriptional profile characterized by progression towards exhaustion and waning signature of cell activation during chronic stimulation *in vitro*.

Some genes upregulated in *in vivo* T_ex_ were not robustly enriched after chronic stimulation *in vitro*, such as *Ctla4* and *Tox* (Fig. S2E). GO analysis of genes absent from the leading edge highlighted pathways related to cell activation as well as cytokine signaling and inflammation (Fig. S2F), further emphasizing antigen-specific TCR stimulation as the basis of our *in vitro* model of chronic stimulation, rather than cytokine/chemokine signaling, inflammation, or other aspects that contribute to development of *in vivo* T_ex_. These observations were supported by low expression of *Tox* mRNA observed after chronic stimulation *in vitro* (Fig. S2G). Furthermore, although TOX protein expression increased marginally after *in vitro* chronic stimulation compared to acute stimulation, induction of TOX in *in vitro* chronically stimulated P14 cells was modest compared to that of *in vivo* T_ex_ (Fig. S2H). Low expression of TOX in this and other previously published *in vitro* models of T_ex_ (*17, 18*) indicates that, although TOX may be required for T_ex_ to form and persist *in vivo*, there are likely features of T_ex_ are less dependent on high and sustained TOX expression that can be captured using *in vitro* models. Thus, these GSEA and GO analyses suggest that chronic stimulation *in vitro* was sufficient to recreate some aspects of transcriptional programs distinctly upregulated in *in vivo* T_ex_, such as IR expression and negative regulation of cell activation, but not others, such as inflammatory signaling and high and sustained expression of TOX.

GSEA also revealed a pattern of genes downregulated in T_ex_ that were likewise significantly negatively enriched in *in vitro* chronically stimulated P14 cells (Fig. 2E). Leading edge genes included *Tcf7*, *Il7r*, *Sell*, *Lef1*, *Ccr7*, and other known T_n_- and T_mem_-associated genes (Fig. S2I). GO analysis of these leading edge genes revealed pathways involved in cell-cell adhesion, WNT, MAPK, ion, and sphingolipid signaling, confirming that *in vitro* chronic stimulation significantly recapitulates transcriptional downregulation of programs associated with quiescence and memory. In total, these GSEA results suggest that, whereas this *in vitro* model of chronic stimulation partially captures transcriptional programming upregulated in *in vivo* T_ex_, this model appeared to broadly capture the transcriptional programs distinctly downregulated in T_ex_.

To compare transcriptional profiles of *in vitro* chronically and acutely stimulated CD8 T cells to those of *in vivo* T_n_, T_eff_, T_mem_, and T_ex_, we computationally merged our RNA-seq data set with a previously published data set of *in vivo* generated antigen-specific CD8 T cells (*37*) and visualized the results by PCA. (Note that differences between T_n_ populations represent naïve P14 cells versus polyclonal naïve CD8 T cells.) PCA revealed co-localization of acutely and chronically stimulated P14 cells at d4 of *in vitro* culture (Fig. 2F), consistent with earlier data (Fig. S2A). Both P14 populations at d4 also co-localized with *in vivo* T_eff_ (Fig. 2F), consistent with a dominant signature of T cell activation at this early time point *in vitro*. By day 7, however, acutely and chronically stimulated P14 cells transcriptionally diverged to occupy distinct principal component space. Whereas *in vitro* acutely stimulated P14 cells at d7 co-localized with *in vivo* T_mem_, *in vitro* chronically stimulated P14 cells at d7 overlapped nearly completely with *in vivo* T_ex_ (Fig. 2F). These data suggest that, during the first 4 days of *in vitro* culture, acute and chronic stimulation drove similar transcriptional programming consistent with programs of *in vivo* T_eff_. However, the absence of further stimulation in acute conditions *in vitro* allowed P14 cells to adopt T_mem_-like transcriptional programming. In contrast, continued chronic antigenic stimulation *in vitro* induced T_ex_-like transcriptional changes.

We next sought to evaluate transcriptional programs induced by acute and chronic stimulation *in vitro* in the context of known T_eff_, T_mem_, and T_ex_ biology. We therefore evaluated expression of key effector-, memory- and exhaustion-associated genes in *in vitro* chronic and acutely stimulated P14 cells (Fig. 2G). By d7, *in vitro* acutely stimulated P14 cells highly expressed many genes associated with T_mem_ (*Bcl6*, *Tcf7*, *Sell*, *Lef1*, *Slamf6*, *Cxcr5*, *Cd27*, and *Ccr7*), reflecting the overlap in principal component space described above (Fig. 2F). In contrast, by d7, chronically stimulated P14 cells upregulated some T_ex_-associated genes (*Pdcd1*, *Havcr2*, *Entpd1*, *Lag3*, *Cd244*, and *Tigit*) but lacked robust expression of others (*Cd160*, *Batf*, *Ctla4, Tox, Nfatc1*). Chronic stimulation also induced expression of some T_eff_-associated genes (*Klrg1*, *Tnf*, *Ifng*, *Eomes*, *Prf1*, *Gzmb*), consistent with increased transcription of T_eff_-related genes despite reduced protein expression by *in vivo* T_ex_. Overall, these data indicate that acute stimulation *in vitro* was sufficient to induce expression of many known T_mem_-related genes, whereas chronic stimulation *in vitro* induced expression of known activation- and T_ex_-related genes. Furthermore, these results suggest that chronic antigenic stimulation *in vitro* was sufficient to induce many T_ex_-associated transcriptional changes but additional stimuli may be necessary to fully induce and/or sustain certain pathways, including the TOX-NFAT axis.

*In vivo* T_ex_ are heterogeneous and consist of biologically distinct subsets, including progenitor (or stem-like), intermediate (T_eff_-like), and terminal populations (*24, 26, 33, 42, 43*). Progenitor T_ex_ express the TF TCF1 and provide the long-term proliferative reserve needed to give rise to intermediate and terminally differentiated subsets of T_ex_ (*24, 27-29*). Progenitor T_ex_ also provide the proliferative burst necessary for response to checkpoint blockade (*24, 27, 29*). Intermediate T_ex_ re-acquire some effector activity and give rise to terminal T_ex_ that are often found in non-lymphoid tissues and tumors (*33, 44*). To investigate whether *in vitro* chronically stimulated P14 cells recapitulated the biology of any of these T_ex_ subsets, we used gene set variation analysis (GSVA) (*45*) to quantify enrichment of transcriptional signatures of progenitor and terminal T_ex_ subsets (*28*) in *in vitro* chronically and acutely stimulated P14 cells. We also included signatures from major CD8 T cell states, such as T_n_, T_eff_, and T_mem_, as well as a signature of bulk T_ex_ which contained progenitor, intermediate, and terminal populations (*41*). Whereas *in vitro* chronically stimulated P14 cells enriched for signatures of T_ex_ and T_eff_ (Fig. 2H), *in vitro* acutely stimulated P14 cells enriched for T_n_ and T_mem_ gene signatures. Furthermore, *in vitro* chronically stimulated P14 cells enriched for a signature of terminal T_ex_, even more strongly than for the signature of bulk T_ex_, and lacked enrichment for a signature of progenitor T_ex_. These data are also consistent with the absence of genes involved in stem and progenitor biology in *in vitro* chronically stimulated P14 cells, such as *Tcf7*, *Lef1*, *Slamf6*, and *Cxcr5* (Fig. 2G). This GSVA data suggests that chronic stimulation *in vitro* may induce a transcriptional profile more representative of terminally differentiated T_ex_.

Progenitor and terminal subsets of T_ex_ are defined not only by differential transcriptional signatures but also by differences in phenotype and function. For example, progenitor and terminal subsets of T_ex_ are associated with discrete TF hallmarks: whereas TCF1 expression is associated with progenitor T_ex_ (*27, 28, 46, 47*), expression of Eomes is associated with terminally differentiated T_ex_ subsets (*26*). We therefore sought to evaluate whether the terminally differentiated transcriptional signature of *in vitro* chronically stimulated P14 cells was associated with phenotypic or functional differences. Indeed, *in vitro* chronically stimulated P14 cells and *in vivo* T_ex_, but not *in vitro* acutely stimulated P14 cells, displayed high co-expression of PD-1 and Eomes (Fig. S2J), consistent with terminal T_ex_ differentiation (*26*). Similarly, whereas bulk *in vivo* T_ex_ had both TCF1+ progenitor and TCF1-terminal populations, TCF1 expression was almost entirely absent in P14 cells after *in vitro* chronic stimulation (Fig. S2K), in line with prior work linking chronic stimulation *in vitro* to *Tcf7* promoter methylation and gene silencing (*17*). Additionally, chronic stimulation *in vitro* led to increased production of Granzyme B compared to bulk *in vivo* T_ex_ (Fig. S2L). This observation aligns with recent studies documenting increased cytotoxic potential in terminal T_ex_ compared to other T_ex_ subsets (*33, 44*). Taken together, these TF expression patterns and increased cytotoxic potential confirm earlier transcriptional data indicating that chronic stimulation *in vitro* induces a cell state with similarities to the terminal subset of *in vivo* T_ex_.

### *In vitro* chronically stimulated CD8 T cells develop epigenetic signatures of T_eff_ and T_ex_

Recent studies have shown that T_ex_ develop an epigenetic landscape distinct from T_eff_ or T_mem_ (*37, 38, 48*). To assess global epigenetic changes induced by acute versus chronic stimulation *in vitro*, we analyzed P14 cells by ATAC-seq on days 0, 4, and 7 of *in vitro* culture. By number of differentially accessible chromatin regions (DACRs), the most substantial differences were observed between naïve (d0) and *in vitro* stimulated P14 cells at d4 (∼30,000 DACRs) regardless of stimulation condition (Fig. S3A), indicating that the most robust epigenetic changes occurred upon the transition from naïve to activated CD8 T cell. Whereas acute and chronically stimulated P14 cells only differed by 547 DACRs at d4 of *in vitro* culture, this number grew to 12160 by d7 (Fig. S3A), suggesting that acute and chronic stimulation induced similar early chromatin accessibility programs that diverged significantly by d7. By PCA, acutely and chronically stimulated P14 cells initially co-localized at d4 but later diverged to occupy distinct principal component space by d7 (Fig. S3B). Collectively, these epigenetic data reveal distinct overall trajectories of differentiation associated with chronic or acute stimulation *in vitro*, consistent with the transcriptional trajectories observed earlier.

To understand how the global epigenetic programs of *in vitro* chronically and acutely stimulated P14 cells compare to those of *in vivo* T_eff_, T_mem_, and T_ex_, we computationally merged our *in vitro* ATAC-seq data set with a previously published ATAC-seq data set of *in vivo* generated CD8 T cell populations (*48*) and visualized the results by PCA (Fig. 3A). P14 cells at d4 of *in vitro* chronic and acute stimulation clustered closely with *in vivo* T_ex_ in principal component space, perhaps reflecting activation-associated chromatin accessibility. However, by d7, acutely stimulated P14 cells co-localized with *in vivo* T_mem_. Chronically stimulated P14 cells at d7 instead remained co-localized near *in vivo* T_ex_ in principal component space, indicating chromatin accessibility at many T_ex_-specific loci after chronic stimulation *in vitro*. Indeed, accessibility at the −23.8kb enhancer in the *Pdcd1* locus, previously described as uniquely accessible in *in vivo* T_ex_ with downstream functional effects on PD-1 expression (*38*), was observed only in *in vitro* chronically stimulated P14 cells and *in vivo* T_ex_ (Fig. 3B) but not in *in vitro* acutely stimulated P14 cells. Chronically stimulated P14 cells also co-localized in principal component space with *in vivo* T_eff_ (Fig. 3A), suggesting that chronic stimulation *in vitro* generated a chromatin accessibility profile with characteristics of both T_eff_ and T_ex_.

**Fig. 3.**
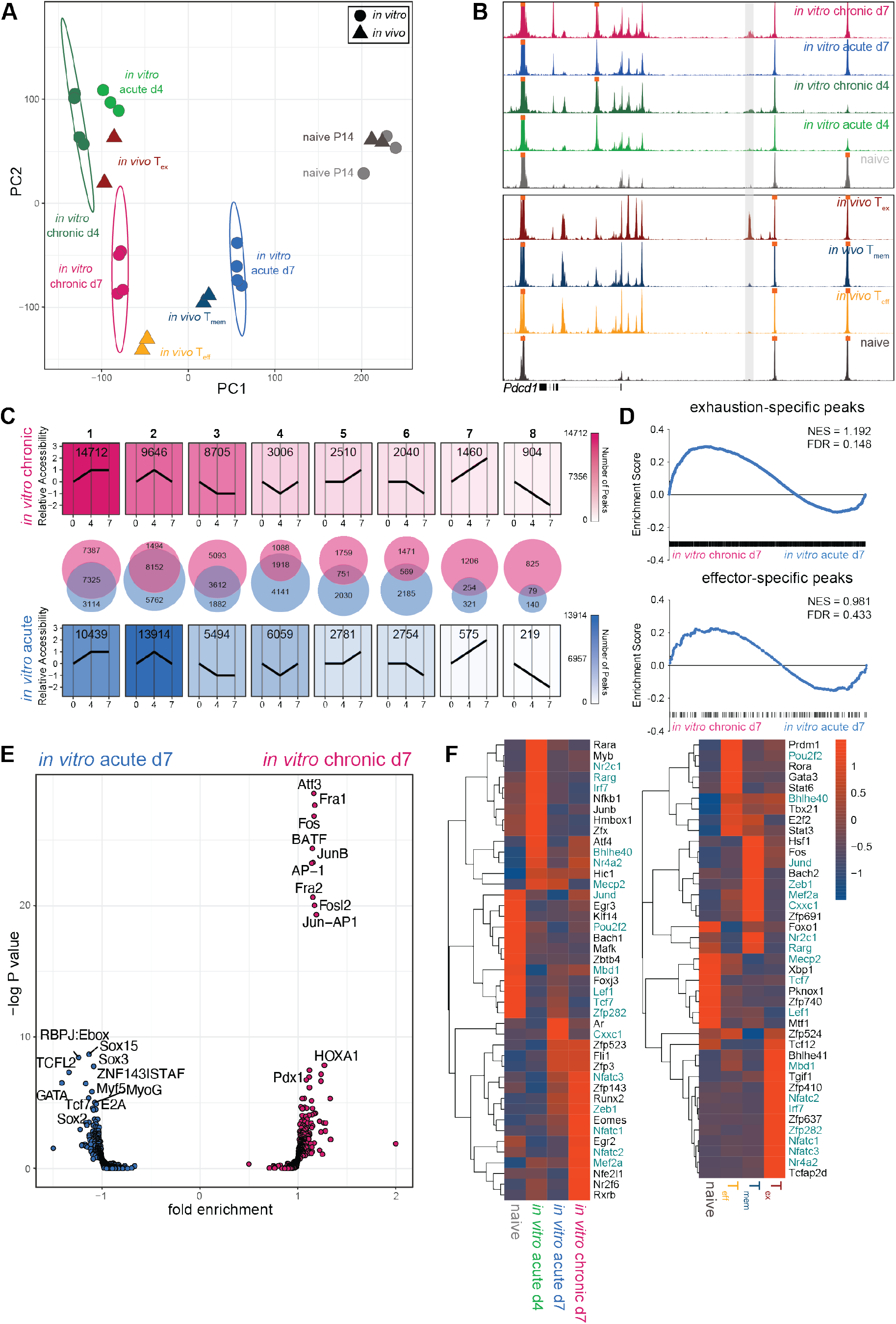
*In vitro* chronically stimulated P14 cells develop epigenetic signatures of T_eff_ and T_ex_. **(A)** PCA of ATAC-seq data from *in vitro* chronically and acutely stimulated P14 cells and previously published *in vivo* CD8 T cell subsets (*48*). **(B)** ATAC-seq signal tracks for *in vitro* chronically and acutely stimulated P14 cells and previously published *in vivo* CD8 T cell subsets (*48*); −23.1kb enhancer of *Pdcd1* locus highlighted in gray. **(C)** Trajectory analysis (see Methods) of chromatin accessibility patterns during *in vitro* chronic or acute stimulation. **(D)** Peak set enrichment analysis (PSEA) of T_ex_- and T_eff_-specific ACRs in *in vitro* chronically and acutely stimulated P14 cells. **(E)** Transcription factor (TF) motif accessibility in differential ACRs between *in vitro* chronically and acutely stimulated P14 cells. **(F)** Rank calculated via Taiji analysis in [left] *in vitro* chronically and acutely stimulated P14 cells and [right] previously published *in vivo* CD8 T cell subsets (*37*), filtered by mean>0.0001 and fold change>5. Heat scale indicates z-score of rank as calculated by PageRank algorithm. Colored text indicates shared TFs between *in vitro* and *in vivo* analyses.

To define the temporal epigenetic changes induced by chronic and acute stimulation *in vitro*, we manually binned individual accessible chromatin regions (ACRs) into patterns based on relative accessibility over the course of *in vitro* culture (Fig. 3C). During chronic stimulation *in vitro*, the most frequent pattern was an increase in chromatin accessibility between d0 and d4, followed by stable maintenance from d4 to d7 (pattern 1). This most frequent pattern indicates that chronic stimulation *in vitro* preserved a majority of ACRs opened during the transition from naïve to activated cell. In contrast, the most common pattern during *in vitro* acute stimulation was a transient increase from d0 to d4 followed by a return to baseline by d7 (pattern 2). This observation is consistent with the notion that, in the absence of further TCR engagement *in vitro*, P14 cells preferentially close many ACRs that became transiently accessible during activation, returning to a state of relative epigenetic inaccessibility. Many ACRs binned into pattern 2 were shared between chronic and acute stimulation *in vitro* (8152 shared ACRs), suggesting some common epigenetic patterning of transient accessibility associated with initial activation. ACRs that progressively decreased in accessibility over *in vitro* culture (pattern 8) represented the least common pattern in both acute and chronic stimulation, followed by ACRs that progressively increased in accessibility (pattern 7). Patterns 7 and 8 were also characterized by the least amount of overlap between *in vitro* chronic and acute stimulation (254 and 79 shared ACRs, respectively), indicating divergent epigenetic landscapes and biology preferentially associated with either chronic or acute stimulation *in vitro*. Overall, these data suggest that, unlike acutely stimulated cells, P14 cells undergoing *in vitro* chronic stimulation induced epigenetic changes consistent with some aspects of T_ex_, including a prominent pattern of sustained changes in accessibility.

To compare chromatin accessibility profiles of *in vitro* chronically and acutely stimulated P14 cells to *in vivo* T_n_, T_eff_, T_mem_, and T_ex_, we performed peak set enrichment analysis (PSEA) (*49*) using signatures of subset-specific ACRs, or “peaks,” curated from a previously published data set (*48*). PSEA revealed enrichment for T_mem_-specific peaks after acute stimulation *in vitro* (Fig. S3C), consistent with shared epigenetic programming of quiescence. Conversely, PSEA of *in vitro* chronically stimulated P14 cells revealed enrichment for T_ex_-specific ACRs (Fig. 3D). Enrichment for T_n_-(Fig. S3C) and T_eff_-specific peaks (Fig. 3D) were not significant, suggesting that both *in vitro* acutely and chronically stimulated P14 cells share some ACRs with *in vivo* T_n_ and T_eff_. Collectively, these PSEA results indicate that acute stimulation *in vitro* induced a chromatin accessibility landscape similar to that of T_mem_ whereas chronic stimulation *in vitro* generated an epigenetic profile similar to that of T_ex_.

To investigate how the epigenetic landscape induced by chronic stimulation *in vitro* might affect downstream TF binding, we performed TF motif enrichment analysis in DACRs between *in vitro* chronically stimulated P14 cells and *in vivo* T_ex_. Whereas ACRs unique to *in vitro* chronic stimulation enriched for binding motifs of Fli1 and RUNX family members, NFAT binding motifs predominated ACRs unique to *in vivo* T_ex_ (Fig. S3D). ACRs unique to *in vivo* T_ex_ also enriched for T-box family motifs and NFAT:AP1 composite motifs. Prior work demonstrated that constitutive expression of partnerless NFAT (engineered to be unable to interact with AP1) can induce expression of TOX and other hallmarks of T_ex_ (*50*) and that NFAT is necessary for induction and stabilization of TOX *in vivo* (*49*). The relative absence of monomeric NFAT binding sites in *in vitro* chronically stimulated P14 cells suggests that antigenic stimulation *in vitro* may not be sufficient to remodel chromatin in a manner that accommodates partnerless NFAT binding to the extent of *in vivo* T_ex_. Furthermore, these results suggest that a comparative lack of monomeric NFAT sites in the chromatin landscape induced by chronic stimulation *in vitro* may be associated with only modest induction of TOX expression (Fig. S2H). This analysis identified differential modules of TF binding potential between *in vitro* chronically stimulated P14 cells and *in vivo* T_ex_, providing context for which transcriptional circuits might be accurately modeled *in vitro*.

To further examine TF binding potential downstream of differential chromatin accessibility between *in vitro* acutely and chronically stimulated P14 cells, we conducted TF motif enrichment analysis in DACRs between these two conditions (Fig. 3E). ACRs unique to *in vitro* acutely stimulated P14 cells enriched for TCF family binding motifs, consistent with the role of these TFs in restoring stemness and quiescence following acute stimulation, as well as Sox family binding motifs. Conversely, ACRs unique to chronically stimulated P14 cells enriched for AP1 family binding motifs, including BATF, Fos, Fosl2, AP1, and Jun-AP1 dimer motifs. Strong enrichment for AP1 family member motifs could potentially indicate disruption of canonical Fos/Jun signaling via formation of heterodimers, leading to a reduction in AP1 available to partner with NFAT, as has been implicated by studies describing the role of BATF in T_ex_ (*51*).

To predict TF networks central to differential biology in these *in vitro* models of chronic and acute stimulation, we computationally integrated chromatin accessibility and TF motif data (from ATAC-seq) with gene expression data (from RNA-seq) and applied Taiji PageRank analysis (*52, 53*). Taiji analysis constructs networks of interactions between TFs to infer importance, or rank, based on connectivity within these networks. Taiji scans motif binding sites in ACRs from ATAC-seq, then assigns TF motifs found in enhancer or promoter regions to target genes. Once TFs have been linked to gene targets, Taiji constructs putative TF networks then weights TF nodes in these networks based on gene expression data from RNA-seq. The PageRank algorithm is then applied to determine rank, or importance, based on node weight and connectivity. Highly ranked TFs will have high gene expression as well as binding motifs present in ACRs at loci of multiple other TFs, indicating that their influence extends to multiple transcriptional networks. To contextualize analysis of our *in vitro* data sets within known CD8 T cell biology, we applied Taiji analysis in parallel to a previously published *in vivo* RNA- and ATAC-seq data set (*37*). Taiji analysis identified many of the same ranked TFs in acute and chronic stimulation *in vitro* and *in vivo* (Fig. 3F). For example, *Lef1* and *Tcf1*, TFs associated with stemness and quiescence, ranked highly in T_n_ as well as *in vitro* acutely stimulated P14 at d7 and *in vivo* T_mem_. *Zeb1* ranked highly in *in vitro* acutely stimulated P14 cells at d7 and *in vivo* T_mem_, confirming a role for this TF in coordinating memory-associated transcriptional networks (*54*). *Zeb1* also ranked highly in *in vitro* chronically stimulated P14 cells at d7, indicating potential activity in T_ex_ biology. Similarly, effector-related TFs, such as *Myb*, *Junb*, *Nfkb1*, and *Fli1*, as well as TFs without previously defined roles in T cell differentiation, such as *Mecp2*, ranked highly in *in vitro* acutely stimulated P14 cells at d4 and *in vivo* T_eff_. Exhaustion-related TFs, such as *Eomes*, *Nr4a2*, and multiple NFAT family members, ranked highly in *in vitro* chronically stimulated P14 cells and *in vivo* T_ex_. Despite the relative lack of enrichment of NFAT binding motifs observed in ACRs of *in vitro* chronically stimulated P14 cells compared to *in vivo* T_ex_ (Fig. S3D), *Nfatc1* and *Nfatc2* emerged as highly ranked TFs via Taiji analysis. This observation suggests that these transcriptional networks have activity and influence during chronic stimulation *in vitro*, though perhaps to a lesser extent than in *in vivo* T_ex_. These data are also consistent with a key role for NFAT circuitry in CD8 T cell exhaustion but suggest that *in vitro* settings may access this biology less efficiently than *in vivo* T_ex_.

Many transcription factors have divergent roles in different cellular and epigenetic contexts. For example, NR4A family members operate downstream of TCR activation but also have non-redundant functions in transcriptional regulation of T_ex_ (*55*). Taiji analysis of *in vitro* generated P14 cells reflected these dual roles in both acute and chronic activation, with *Nr4a2* ranking highly in *in vitro* acutely stimulated P14 cells at d4 and *in vitro* chronically stimulated P14 cells at d7 (Fig. 3F). This observation suggests persistent or preferential use of *Nr4a2* activity in the trajectory toward T_ex_, consistent with *in vivo* observations (*55*). Taiji analysis revealed another key transcriptional circuit centered around *Bhlhe40*. *Bhlhe40* was predicted to have divergent functions in both effector- and exhaustion-associated biology due to rank in *in vitro* acutely stimulated P14 cells at d4 and chronically stimulated P14 cells at d7; *Bhlhe40* also ranked highly in both *in vivo* T_eff_ and T_ex_. As such, *Bhlhe40* could represent a point of convergence for transcriptional control of both T_eff_ and T_ex_ biology. Identification of these TFs suggests that Taiji analysis can reveal TFs with non-overlapping roles in these distinct cellular contexts.

### Pooled CRISPR screening identifies novel transcriptional regulators of CD8 T cell differentiation

To leverage this *in vitro* model as a discovery tool and interrogate regulators of T_ex_, we used a pooled CRISPR screening approach (*56*). We applied the same sgRNA library previously used in *in vivo* CRISPR screening of T_ex_ and T_mem_ (*56, 57*) to facilitate comparison of results from *in vitro* versus *in vivo* CRISPR screening. Using P14 cells from Cas9^+/-^ GFP P14 mice, *in vitro* chronic and acute stimulation proceeded as described above. At ∼24 hours post-activation, Cas9^+/-^ GFP P14 cells were transduced with a pooled sgRNA VEX-expressing retrovirus (RV) library targeting 150 TFs (*56, 57*). Transduced P14 cells were then returned to co-culture under acute or chronic stimulation conditions *in vitro*. VEX+GFP+ P14 cells were sorted and sgRNA cassettes were sequenced at day 2 (baseline) and day 7 of *in vitro* culture (Fig. 4A). Positive and negative selection at the population level were quantified via sequencing.

**Fig. 4.**
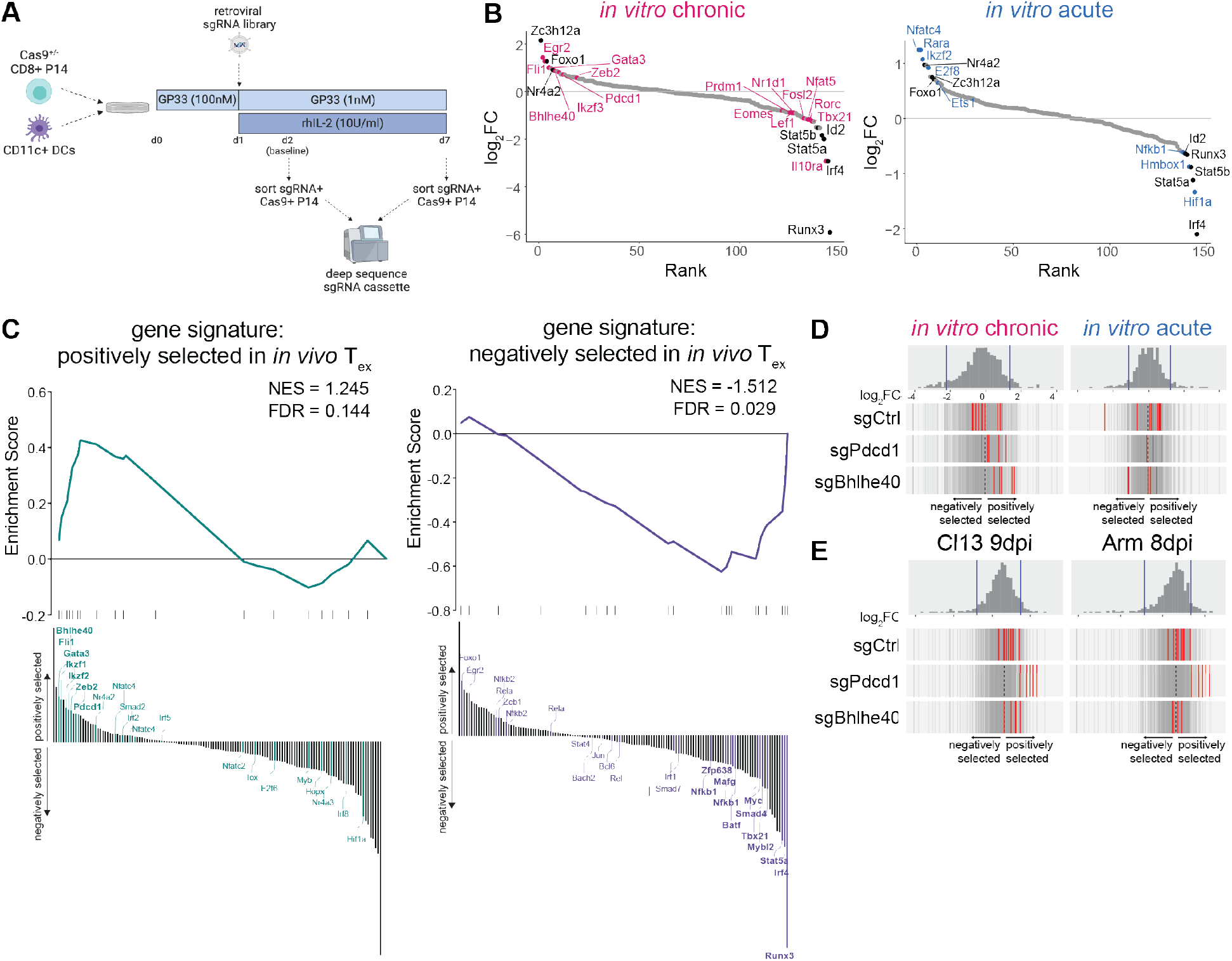
Pooled CRISPR screening in *in vitro* chronically stimulated P14 cells identifies novel transcriptional regulators of CD8 T cell differentiation. **(A)** Experiment schematic of pooled CRISPR screening in *in vitro* chronically stimulated P14 cells. **(B)** Genes targeted by pooled CRISPR screen ranked by magnitude of selection (as quantified by lfc) after acute and chronic stimulation *in vitro*. Colored text indicates hits unique to each condition; black text indicates hits common to both. **(C)** GSEA of gene sets constructed from [left] positively selected and [right] negatively selected sgRNAs from previously published pooled *in vivo* CRISPR screening (*57*). Waterfall plot illustrates rank order of hits in *in vitro* chronically stimulated P14 cells; bold lettering indicates leading edge genes. **(D-E)** Selection of individual sgRNAs against negative control genes, *Pdcd1,* and *Bhlhe40* **(D)** after acute and chronic stimulation *in vitro* and **(E)** from *in vivo* data set (*57*) in PBMC at 9dpi of LCMV-Cl13 and 8dpi of LCMV-Arm. Histogram and vertical gray bars indicate distribution of all sgRNAs; vertical red bars represent indicated sgRNAs. **(B-D)** Representative of two experiments (two technical replicates and 2 biological replicates each).

Hits from *in vitro* pooled CRISPR screening included many TFs with known roles in CD8 T cell biology. For example, sgRNAs negatively selected after both *in vitro* acute and chronic stimulation included TFs involved in CD8 T cell activation and proliferation, such as *Id2*, *Irf4*, *Runx3*, *Stat5a*, and *Stat5b* (Fig. 4B), identifying these as required factors for optimal CD8 T cell differentiation *in vitro*. sgRNAs positively selected after both *in vitro* acute and chronic stimulation included *Foxo1* and *Nr4a2*, consistent with prior literature describing these TFs in restraining initial T cell activation and effector development (*55, 58-60*). This screen also identified targets preferentially associated with chronic stimulation *in vitro*, including several TFs involved in transcriptional regulation of T_ex_. As a positive control for T_ex_-specific biology, sgRNAs against *Pdcd1* were included; as expected, these sgRNAs were among the top positively selected hits after chronic stimulation *in vitro*. Other sgRNAs positively selected after chronic stimulation *in vitro* included those targeting TFs with known roles in activation and effector function, such as *Fli1* and *Zeb2* (*54, 57, 61, 62*), as well as TFs with recently identified roles in regulation of stability and/or maintenance of T_ex_, such as *Gata3* and *Egr2* (*63–65*) (Fig. 4B). sgRNAs negatively selected after chronic stimulation *in vitro*, such as *Eomes*, *Tbx21*, *Prdm1*, and *Nfat5*, reflected transcriptional targets previously implicated in T_ex_ (*26, 49, 50, 66-69*). Identification of these TFs with known roles in CD8 T cell differentiation and function confirms that these *in vitro* models of acute and chronic stimulation can not only accommodate high-throughput assays such as pooled CRISPR screening, but also recapitulate known biology for cell types of interest.

To benchmark this *in vitro* CRISPR screening approach against *in vivo* T_ex_, we next compared *in vitro* CRISPR screen data to a previously published CRISPR screen in *in vivo* T_ex_ that used the same sgRNA library (*57*). To quantify overlap between hits from CRISPR screening *in vitro* and *in vivo*, we generated gene sets from positively or negatively selected hits from CRISPR screening of *in vivo* T_ex_ and used GSEA to evaluate enrichment of these gene sets *in vitro*. GSEA revealed significant enrichment of the positively selected *in vivo* T_ex_ gene set in hits positively selected after chronic stimulation *in vitro* (Fig. 4C). Leading edge genes included TFs strongly implicated in T_ex_ biology *in vitro* and *in vivo*, such as *Bhlhe40*, *Fli1*, *Gata3*, *Ikzf1/2*, and *Zeb2*. Conversely, positively selected hits *in vivo* that did not appear in the leading edge of overlap with *in vitro* T_ex_ included *Tox*, *Nfatc2*, and *Nfatc4*, highlighting the relative dearth of the TOX/NFAT signaling axis *in vitro*. *Hif1a*, though strongly positively selected in *in vivo* T_ex_, was negatively selected after chronic stimulation *in vitro*, pointing to differential regulation of hypoxia-related transcriptional signaling, potentially due to non-hypoxic conditions *in vitro*. GSEA also showed significant enrichment of the negatively selected *in vivo* T_ex_ gene set in hits negatively selected after chronic stimulation *in vitro* (Fig. 4C). Leading edge genes included TFs associated with activation and effector programs, such as *Runx3*, *Irf4*, *Tbx21*, *Myc*, *Batf* (Fig. 4C). Negatively selected hits *in vivo* that were not replicated *in vitro* included TFs such as *Foxo1*, *Egr2*, and *Bcl6*, reflecting the relative absence of transcriptional networks associated with stem-like T_ex_ progenitors during chronic stimulation *in vitro*. This analysis comparing results from CRISPR screening in both chronic stimulation *in vitro* and *in vivo* T_ex_ highlight the ability of this *in vitro* model of chronic stimulation to recapitulate transcriptional networks that coordinate key aspects of T_ex_ biology. Conversely, analysis of differentially selected TFs between these two models could be useful for parsing pathways responsive to other environmental factors unique to *in vivo* T_ex_, such as NFAT signaling, inflammation, or hypoxia.

After benchmarking pooled CRISPR screening in our *in vitro* model of chronic stimulation to previously published *in vivo* CRISPR screening with the same library, we probed our *in vitro* dataset further to identify novel regulators of T_ex_. BHLHE40 was of particular interest because it was previously identified via Taiji analysis as a highly ranked TF in both *in vitro* chronically stimulated P14 cells and *in vivo* T_ex_ (Fig. 3F). *Bhlhe40* was also among the most robustly enriched leading edge genes positively selected in CRISPR screening of chronic stimulation *in vitro* and *in vivo* T_ex_ (Fig. 4C). All sgRNAs against *Bhlhe40* were positively selected after chronic stimulation *in vitro*, with some displaying stronger selection than guides against *Pdcd1* (Fig. 4D). Conversely, sgRNAs against *Bhlhe40* were not positively selected after acute stimulation *in vitro*, indicating biology biased toward chronic stimulation. *Bhlhe40* also displayed positive selection in *in vivo* CRISPR screening in chronic (Cl13) but not acute infection (Arm) (Fig. 4E), confirming T_ex_-specific biology *in vivo*.

### BHLHE40 is a transcriptional regulator of T_ex_ differentiation

Pooled CRISPR screening and Taiji analysis of the *in vitro* model of chronic stimulation both identified BHLHE40 as a potential regulator of T_ex_ and T_eff_ biology. BHLHE40 has been previously implicated in several aspects of T cell biology, including regulation of CD4 T cell lineage commitment and differentiation (*70, 71*) and maintenance of stemness in CD4 and CD8 tumor-infiltrating lymphocytes (TIL) as well as tissue-resident memory CD8 T cells (*72, 73*). Moreover, BHLHE40 has recently been implicated in the response of CD8 TIL to checkpoint blockade therapy (*74*). However, it is unclear whether BHLHE40 functions differently in the formation of T_ex_ versus T_eff_ and, if so, how BHLHE40 might regulate establishment and/or maintenance of T_ex_. Furthermore, T_ex_ biology includes a hierarchy of subsets linked to developmental lineage and differential response to checkpoint blockade; it is unclear where in this hierarchy BHLHE40 impacts T_ex_.

To determine whether BHLHE40 has the potential to differentially regulate T_eff_ and T_ex_, we first used Taiji analysis to construct TF networks downstream of BHLHE40 in *in vitro* acutely stimulated P14 cells at d4 versus *in vitro* chronically stimulated P14 cells at d7. Some TFs were common to both settings; however, most TFs in downstream networks regulated by BHLHE40 were found in only one condition (Fig. S4A). For example, *Bcl6*, *Rxra*, *Runx1*, and several KLF family members preferentially connected downstream of BHLHE40 after acute stimulation *in vitro*. In contrast, after chronic stimulation *in vitro*, BHLHE40 connected to a set of Ets family TFs. These observations suggest cell context-specific transcriptional regulation by BHLHE40. We also performed TF co-occurrence analysis to identify TFs that might bind chromatin in tandem with BHLHE40. We found many TFs for which binding motifs co-occur within <100bp proximity of binding motifs of BHLHE40, indicating potential of these TFs to co-bind and co-regulate transcription with BHLHE40 (Fig. S4B); for example, binding motifs for BHLHE40 and BCL6 occur in close proximity at the *Runx1* promoter (Fig. S4C). Because RUNX1 is capable of partially antagonizing RUNX3, which, in turn, promotes effector responses (*57*), our data suggests that co-repression of RUNX1 by BHLHE40 and BCL6 could allow more effective RUNX3-driven T_eff_ circuitry in intermediate T_ex_. Taken together, these results highlight ways in which BHLHE40 may cooperate with other known TFs to regulate CD8 T cell differentiation.

To interrogate the role of BHLHE40 in CD8 T cell differentiation, we employed a loss-of-function approach using shRNA knockdown (KD). We first tested this approach in our *in vitro* model of chronic stimulation (Fig. S5A). shRNA KD led to an 82% reduction in gene expression of *Bhlhe40* (Fig. S5B). In *in vitro* chronically stimulated P14 cells, *Bhlhe40* KD resulted in a decrease in percent expression and MFI of multiple IRs, including PD-1, TIM3, and 2B4 (Fig. S5C), suggesting that BHLHE40 may promote IR expression and providing further support for a potential role for BHLHE40 in CD8 T cells during chronic stimulation.

Whereas our *in vitro* model of chronic stimulation allowed us to screen many TFs in parallel and identify genes of interest, including *Bhlhe40*, we next used the LCMV model to test if and how BHLHE40 regulates CD8 T cell differentiation and function *in vivo*. We thus employed shRNA KD of *Bhlhe40* during viral infection with the acutely-resolving Armstrong strain (Arm) or chronic clone 13 strain (Cl13) of LCMV (Fig. 5A). Congenically distinct P14 cells were transduced with an RV expressing either a control shRNA or shRNA against *Bhlhe40* and adoptively co-transferred at a 1:1 ratio (Fig. S5D) into Arm- or Cl13-infected recipient mice as described (*75*). At 8 days post infection (dpi), *Bhlhe40* KD resulted in a competitive numerical advantage in Cl13 infection (Fig. S5E) but not in Arm (Fig. S5F). However, by 22-31dpi of Cl13, this advantage of *Bhlhe40* KD increased to ∼70-80% of transferred P14 cells (Fig. 5B). These data confirm results from CRISPR screening in the *in vitro* model in which absence of *Bhlhe40* led to a proliferation and/or survival advantage in settings of chronic but not acute stimulation (Fig. 4B,D).

**Fig. 5.**
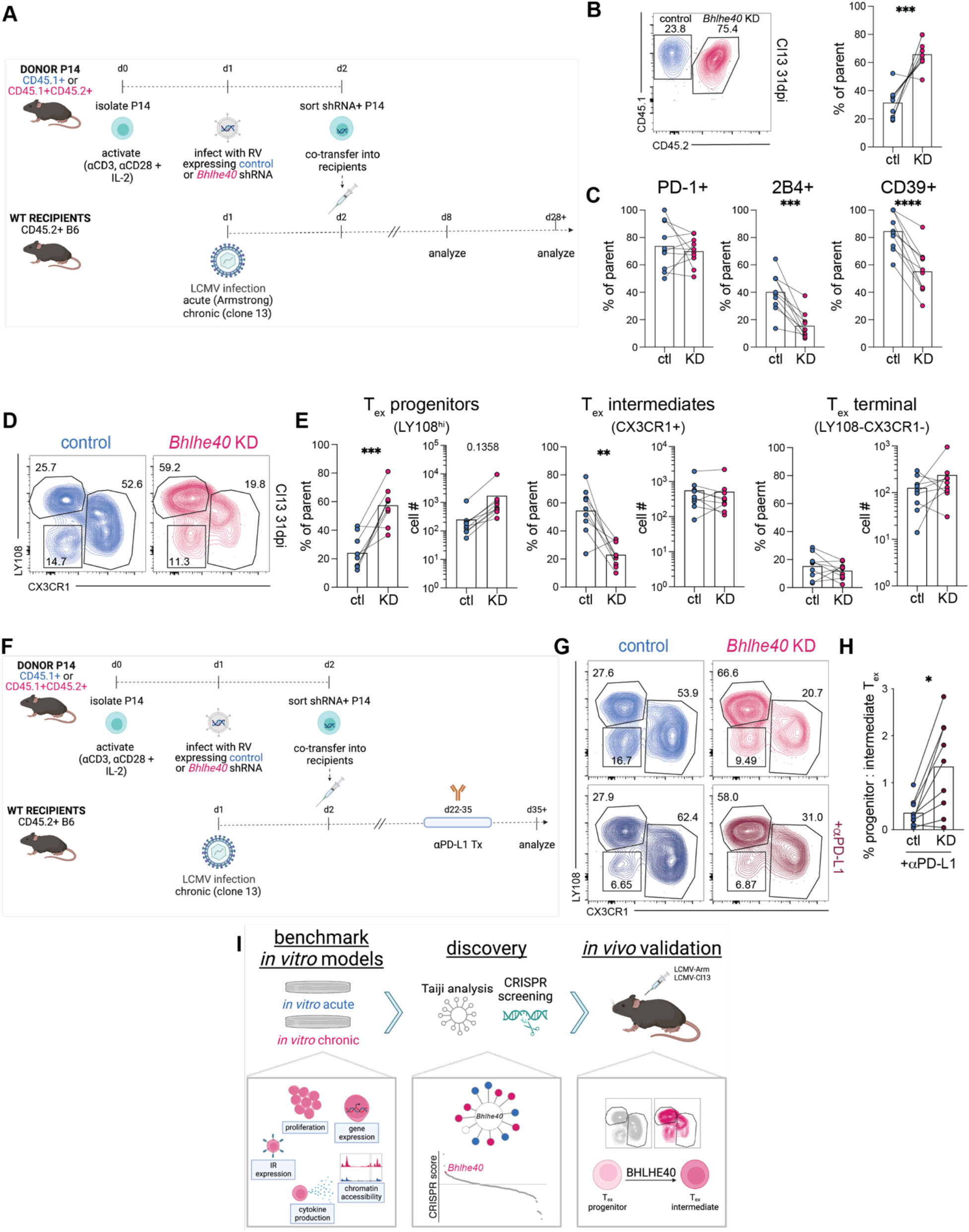
BHLHE40 is a transcriptional regulator of T_ex_ differentiation. **(A)** Experiment schematic of adoptive co-transfer of control and *Bhlhe40* knockdown (KD) P14 cells into LCMV Arm- or Cl13-infected recipient mice. **(B)** [left] Concatenated (n=10) flow cytometry plot and [right] summary data of frequency of control and *Bhlhe40* KD P14 cells (gated on GFP+ CD44^hi^ CD8+ live singlets) at 31dpi of LCMV-Cl13. **(C)** Percent expression of IRs in control and *Bhlhe40* KD P14 cells at 31dpi of LCMV-Cl13. **(D)** Concatenated (n=10) flow cytometry plots of frequency of T_ex_ subsets in control and *Bhlhe40* KD P14 cells at 31dpi of LCMV-Cl13. **(E)** Frequency and total number of progenitor, intermediate, and terminal T_ex_ in control and *Bhlhe40* KD P14 cells at 31dpi of LCMV-Cl13. **(F)** Experiment schematic of adoptive co-transfer of control and *Bhlhe40* KD P14 cells into LCMV Cl13-infected recipient mice, with αPD-L1 treatment or vehicle control between 22-35dpi. **(G)** Concatenated (n=10) flow cytometry plots of frequency of T_ex_ subsets in control and *Bhlhe40* KD P14 cells at 37dpi of LCMV-Cl13, after treatment with vehicle control [top] or αPD-L1 [bottom] from 22-35dpi. **(H)** Ratio of progenitor to intermediate T_ex_ in control and *Bhlhe40* KD P14 cells after αPD-L1 treatment. **(I)** Schematic detailing discovery pipeline. **(B-G)** n=10 mice, representative of 3 experiments. Significance calculated by paired two-tailed t test; **\***p<0.05, **\*\***p<0.01, **\*\*\***p<0.001, **\*\*\*\***p<0.0001. **(B,D,G)** Numbers in flow cytometry plots indicate percentage of parent population within each gate.

Next, to investigate whether BHLHE40 impacts phenotype and function early after infection, we analyzed subset distribution and differentiation states of T_eff_ in control or *Bhlhe40* shRNA-transduced P14 cells in Arm and Cl13. At 7dpi of Arm infection, shRNA KD of *Bhlhe40* led to significantly decreased proportions and numbers of KLRG1+ T_eff_ (Fig. S5G). Furthermore, *Bhlhe40* KD significantly reduced per-cell protein expression (MFI) of KLRG1 (Fig. S5H). At 7dpi of Cl13 infection, before the formation of mature T_ex_, *Bhlhe40* KD also resulted in decreased proportions of KLRG1+ T_eff_-like cells (Fig. S5I). These data support the idea that *Bhlhe40* regulates terminal T_eff_ differentiation during the first week of infection. However, production of TNF and IFNγ was not impacted by *Bhlhe40* KD (Fig. S5J), suggesting that BHLHE40 controls differentiation state rather than effector function.

We then assessed impact of loss of *Bhlhe40* on T_ex_ differentiation at a late time point in Cl13 infection after which exhaustion had been established (*49, 76*). By 31dpi of Cl13, shRNA KD of *Bhlhe40* reduced percent expression and MFI of some IRs, including 2B4 and CD39, but not others, such as PD-1 (Fig. 5C, Fig. S5K,L), confirming earlier observations from the *in vitro* model of chronic stimulation (Fig. S5C). Furthermore, because IR co-expression is associated with progression toward terminal T_ex_ (*26, 32, 33*), these results suggest that Bhlhe40 may promote terminal differentiation of T_ex_.

We next examined the effect of *Bhlhe40* KD on the differentiation hierarchy between subsets of T_ex_ using expression of LY108 and CX3CR1: high expression of LY108 (a surrogate of TCF1 expression) (*33*) distinguishes the progenitor T_ex_ subset, CX3CR1 expression identifies the intermediate and more effector-like T_ex_ subset, and the LY108-CX3CR1-population forms the terminally differentiated T_ex_ subset (*42, 43*). shRNA KD of *Bhlhe40* resulted in an increase of the T_ex_ progenitor subset: whereas roughly 25% of control-transduced P14 cells were progenitor T_ex_, this population increased to almost 60% after *Bhlhe40* KD (Fig. 5D). This increase was seen by percent and absolute number (Fig. 5E). In contrast, loss of *Bhlhe40* led to decreased proportions of the T_ex_ intermediate subset, which represented ∼50% of control-transduced P14 cells but only ∼20% after *Bhlhe40* KD (Fig. 5D). However, the total number of intermediate T_ex_ was similar due to the overall numerical advantage of *Bhlhe40* KD P14 cells (Fig. 5E). *Bhlhe40* KD did not affect subset distribution of terminal T_ex_ (Fig. 5D,E). Because *Bhlhe40* KD resulted in accumulation of T_ex_ progenitors at the expense of T_ex_ intermediates, these data point to a role for *Bhlhe40* in controlling conversion from T_ex_ progenitors into effector-like T_ex_ intermediates. We also examined the impact of *Bhlhe40* gain-of-function via retroviral overexpression (Fig. S5M) on T_ex_ differentiation and found the opposite effect: overexpression of *Bhlhe40* promotes differentiation of T_ex_ intermediates at the expense of both progenitor and terminal T_ex_ (Fig. S5N,O).

Response to immunotherapy, such as PD-1 blockade, hinges on differentiation of T_ex_ progenitors into T_ex_ intermediates (*24, 27, 29, 42, 43*); thus, these results also implicate BHLHE40 in regulating this crucial checkpoint in the response to immunotherapy. To evaluate whether the accumulation of T_ex_ progenitors associated with *Bhlhe40* KD would prove advantageous in synergy with PD-1 pathway blockade, we deployed *Bhlhe40* KD in combination with αPD-L1 treatment during Cl13 infection (Fig. 5F). Regardless of αPD-L1 treatment, *Bhlhe40* KD led to increased numbers and percentages of T_ex_ progenitors (Fig. 5G, S5P). In combination with *Bhlhe40* KD, αPD-L1 treatment resulted in increased percentages of CX3CR1+ intermediates (Fig. 5G, S5P) but this increase was not significant for absolute number of cells. Although PD-1 pathway blockade increased the proportion of T_ex_ intermediates, the accumulation of T_ex_ progenitors created by *Bhlhe40* KD remained intact: whereas the ratio of progenitor to intermediate T_ex_ in control shRNA plus αPD-L1 was roughly 0.5:1, this ratio was inverted to approximately 1.5:1 in *Bhlhe40* KD plus αPD-L1 conditions (Fig. 5H). Overall, these data suggest that, despite the ability of PD-1 pathway blockade to induce a burst of proliferation and differentiation of T_ex_ progenitors into T_ex_ intermediate cells, blocking PD-1 signaling is largely insufficient to override transcriptional reprogramming of this differentiation checkpoint maintained by shRNA KD of *Bhlhe40*.

Overall, these data suggest that *in vitro* screening using a model that captures a subset of T_ex_ biology can identify targets that have specific roles at a distinct stage of *in vivo* T_ex_ differentiation. This approach of chronic stimulation *in vitro*, from benchmarking against known T_ex_ biology to incorporation of pooled CRISPR screening to *in vivo* validation, represents a discovery pipeline that can be applied to identifying additional novel regulators of T_ex_ biology (Fig. 5I).

## DISCUSSION

The inability to durably reverse CD8 T cell exhaustion remains a major barrier to the treatment of cancer and chronic viral infection. A better understanding of the molecular cues that induce and maintain exhaustion will enable development of more effective therapeutics to prevent or reverse this state of CD8 T cell differentiation. *In vivo* models have engendered foundational knowledge about T_ex_. However, *in vivo* models generate low numbers of T_ex_ and are time-consuming, costly, and difficult to scale. These systems are thus less than ideal for generating large amounts of biological material for high-throughput screening techniques or approaches such as proteomics and/or metabolomics. *In vitro* models that can recapitulate many or even some features of T_ex_ are an attractive alternative to *in vivo* models due to their scalability, speed, efficiency, and ability to be customized. Here, we established a model of *in vitro* chronic antigenic stimulation and comprehensively defined the functional, phenotypic, transcriptional, and epigenetic features of this model in comparison to known *in vivo* T_ex_ biology. Understanding which features of exhaustion can be recapitulated and interrogated *in vitro* allowed us to then use this *in vitro* model to perform high-throughput CRISPR-based screening to identify novel transcriptional regulators of T_ex_.

To induce features of T_ex_ *in vitro*, we used chronic antigenic stimulation as the primary driver of exhaustion biology. We opted to mimic physiological TCR engagement by stimulating P14 cells with cognate peptide presented by professional antigen-presenting cells (APCs). Our goal was to induce high antigenic stimulation with peptide presented by a professional APC, then reduce stimulation strength ∼100-fold, consistent with the physiological dynamics of viral or antigen load in chronic infections (*11*). We also aimed to limit the amount of rest between stimulations to achieve continuous rather than intermittent stimulation. We observed, as have other studies using repeated TCR stimulation to mimic exhaustion *in vitro* (*14, 15, 17–20*), that CD8 T cells displayed decreased proliferative potential, high expression of IRs, and reduced production of effector cytokines after chronic stimulation *in vitro*. Benchmarking these phenotypic, transcriptional, and epigenetic features of our *in vitro* model of chronic stimulation against *in vivo* CD8 T_ex_ revealed where this model recapitulated known features of T_ex_ and where this *in vitro* approach diverged from *in vivo* biology. Many aspects of exhaustion biology were robustly captured by chronic stimulation *in vitro*: for example, *in vitro* chronically stimulated P14 expressed IRs at population and per-cell levels comparable to or even higher than *in vivo* T_ex_. Yet other aspects of *in vivo* biology were incompletely captured *in vitro*, including the NFAT/TOX signaling axis. However, defining which aspects of exhaustion biology were captured using this *in vitro* model provided a foundation to leverage this approach for discovery-based assays downstream.

A notable feature of our *in vitro* model of chronic stimulation is the generation of multiple phenotypic, functional, transcriptional, and epigenetic features of T_ex_ despite relatively low gene and protein expression of TOX. Previous studies have identified TOX as a key transcriptional and epigenetic regulator essential for the formation of T_ex_ (*49, 77–79*). However, because T_ex_ fail to develop *in vivo* in the absence of TOX, interrogating which exhaustion-related pathways function dependently and independently of TOX has been difficult. The failure of T_ex_ to persist in the absence of TOX likely reflects a TOX-dependent T_ex_ program of survival and durability, in addition to epigenetic effects, in the setting of chronic antigen stimulation (*49, 77, 78*). However, lack of knowledge about TOX-independent aspects of the T_ex_ program suggests that such pathways may only be apparent using *in vitro* models. Our *in vitro* model of chronic stimulation, in which many phenotypic, functional, transcriptional, and epigenetic features of T_ex_ are generated in the absence of high and sustained expression of TOX, may represent an opportunity to mechanistically interrogate these T_ex_-related pathways to reveal novel therapeutic targets.

*In vivo* T_ex_ form a proliferative hierarchy consisting of progenitor subsets that differentiate into downstream intermediate and terminal subsets. Chronic stimulation *in vitro* largely induced transcriptional, phenotypic, and functional features that overlapped robustly with those of *in vivo* terminal T_ex_, based on enrichment of a gene signature of terminally differentiated T_ex_, high co-expression of IRs, terminally differentiated TF signature, and increased cytotoxic potential after chronic stimulation *in vitro*. It will be interesting to interrogate whether P14 cells subjected to chronic stimulation *in vitro* first pass through progenitor and intermediate states before undergoing accelerated terminal differentiation or if these cells bypass upstream differentiation steps in favor of directly inducing features of terminal T_ex_. If chronic stimulation *in vitro* is indeed insufficient for establishment and maintenance of progenitor T_ex_, it is likely that the generation of cells with features of progenitor T_ex_ *in vitro* may require soluble cytokine/chemokine cues or even periods of rest or “rescue” to allow for upregulation of stem- and memory-like transcriptional pathways (*20, 80*). Although TCF1 signaling is known to be requisite for development of progenitor T_ex_ *in vivo* (*81*), the cell-extrinsic cues that drive this transcriptional programming remain poorly understood. Future studies could apply gain-of-function screens to *in vitro* modeling of T_ex_ to address this question.

The *in vitro* T_ex_ model described here has generated many insights into CD8 T cell exhaustion; however, no single *in vitro* model will completely capture the complex *in vivo* biology of T_ex_. An advantage of the current work was benchmarking against bona fide *in vivo* T_ex_ generated via the LCMV model of chronic viral infection in which CD8 T cell exhaustion was first defined (*82, 83*). Comparison to *in vivo* T_ex_ identified many ways in which this *in vitro* model recapitulates known exhaustion biology; this benchmarking also identifies ways it does not (e.g. TOX/NFAT signatures). Furthermore, other models of *in vitro* T_ex_ (*14, 19, 20*) may capture slightly different aspects of exhaustion biology. *In vitro* approaches can be tuned to address different features of known *in vivo* biology: for example, by introducing hypoxia or known inflammatory mediators. As the potential use of *in vitro* models of T_ex_, including variations of engineered cell therapies, expands, benchmarking to define the bounds of biology accurately captured will prove useful. *In vitro* T_ex_ models may also have utility in medium- to high-throughput drug or chemical screening, approaches not feasible *in vivo*. Thus, well-annotated *in vitro* T_ex_ models should have considerable applicability due to scalability, ease of use, and ability to enable novel types of investigation into cellular mechanisms of exhaustion.

Another advantage of this and other *in vitro* models of T cell differentiation is the ability to apply high-throughput discovery assays due to increased efficiency, flexibility, and cell yield. We illustrated the utility of this *in vitro* model in combination with pooled CRISPR screening toward the discovery of novel biology. Compared to recently published *in vivo* CRISPR screens (*56, 57, 84*), we performed CRISPR screening in our *in vitro* model of chronic stimulation with relative logistical ease. For example, due to sgRNA library size and cell recovery requirements, *in vivo* CRISPR screening in an adoptive transfer model required pooling of 4-6 donor P14 mice and upwards of 30 recipient mice per time point per infection (i.e. over 120 total mice) (*57*). Using the same sgRNA library, we were able to perform our experiments with 2 mice each and used each P14 mouse as a biological replicate. Additionally, whereas the *in vivo* adoptive transfer model required 12+ hours of flow cytometric sorting (as well as proportional amounts of requisite reagents) to isolate CRISPR-edited cells, we performed sorting for these cells in under 2 hours.

Comparative analysis of transcriptional and epigenetic data from this *in vitro* model of chronic stimulation and *in vivo* T_ex_ enabled interpretation of results from CRISPR screening within the context of which aspects of T_ex_ biology were accurately recapitulated *in vitro*. For example, because the TOX/NFAT signaling axis was transcriptionally underrepresented in our *in vitro* model of chronic stimulation, we anticipated a lack of selection for these TFs during CRISPR screening. Similarly, because we did not incorporate modeling of hypoxia *in vitro*, *Hif1a* did not emerge as a positively selected hit, even though this TF was positively selected during *in vivo* CRISPR screening. However, CRISPR screening in our *in vitro* model of chronic stimulation identified many TFs with known roles in general CD8 T cell biology, including hits relating to T cell activation, such as *Fli1* and *Zeb2*, as well as TFs with known roles in T_ex_, such as *Eomes*, *Tbx21*, and *Prdm1*. CRISPR screening also uncovered recently described TFs or TFs with previously undescribed roles in T_ex_ biology, such as *Egr2*, *Gata3*, and *Bhlhe40*.

Both Taiji analysis and pooled CRISPR screening in our model of *in vitro* chronic stimulation identified BHLHE40, a novel transcriptional regulator of CD8 T cell differentiation. Prior work has shown that BHLHE40 has a role in anti-tumor CD8 T cell immunity (*73*). However, the mechanism of action of BHLHE40 has remained incompletely understood. In this study, we demonstrated via *in vivo* loss- and gain-of-function experiments that BHLHE40 controls a differentiation checkpoint between progenitor and intermediate T_ex_ subsets. Progenitor T_ex_ have garnered attention because of their role in seeding proliferation and differentiation events underlying response to PD-1 pathway blockade (*24, 27, 29*). However, intermediate T_ex_, due to their numerical expansion and upregulation of effector-associated genes, are postulated to be the functional cellular currency necessary to exert therapeutic control over infection or tumor burden upon checkpoint blockade (*42, 43*). Our finding that decreased BHLHE40 activity results in accumulation of progenitor T_ex_ at the expense of intermediate T_ex_ supports a key role for this TF in the transition from progenitor to intermediate T_ex_ and indicates that intact BHLHE40 activity is likely required for formation of intermediate T_ex_. Notably, PD-1 pathway blockade was insufficient to override transcriptional reprogramming by *Bhlhe40* knockdown. These data suggest that unimpaired function of BHLHE40 is also likely required for robust cellular responses to checkpoint blockade by facilitating increased conversion of progenitor to intermediate T_ex_. This observation is consistent with recent work describing decreased efficacy of checkpoint blockade therapy in genetic deficiency of *Bhlhe40* in a transplantable tumor model (*74*). An appropriate balance between progenitor and intermediate subsets is likely required for optimal T_ex_ function, both at steady state in established exhaustion and in response to checkpoint blockade therapy. Our finding that BHLHE40 and, possibly, other TFs can be leveraged to manipulate this balance has powerful therapeutic implications.

From benchmarking our *in vitro* model of chronic stimulation against known *in vivo* T_ex_ to perturbing this model and identifying functionally relevant TFs that regulate T_ex_ biology, we have demonstrated that *in vitro* models can increase our understanding of the molecular mechanisms that underlie T_ex_. This discovery pipeline can ultimately be used to uncover novel actionable pathways that can be exploited towards the therapeutic reversal of T cell exhaustion in chronic viral infection and cancer.

## MATERIALS AND METHODS

### Mice

Mice were maintained in a specific-pathogen-free facility at the University of Pennsylvania (UPenn). Experiments and procedures were performed in accordance with the Institutional Animal Care and Use Committee (IACUC) of UPenn. P14 transgenic mice expressing a TCR specific for the LCMV peptide D^b^GP^33-41^ were bred in-house and backcrossed onto the C57BL/6 background (Charles River); recipient WT C57BL/6 mice were purchased from Charles River. LSL-Cas9-GFP mice were purchased from Jackson Laboratory (JAX) and bred to CD4CRE and P14 mice on the JAX C57BL/6 background (referred to as C9P14). For *in vitro* experiments, mice were between 8-20 weeks of age; for *in vivo* experiments, mice were between 6-8 weeks of age. For all experiments, mice were age- and sex-matched and randomly assigned to experimental groups.

### Adoptive T cell transfer

CD8 T cells were isolated from peripheral blood of donor P14 mice via gradient centrifugation with Histopaque-1083 (Sigma-Aldrich). 500 naïve P14 cells were adoptively transferred intravenously (i.v.) into 6-8-week-old recipient mice 1 day prior to infection. Recipients were of a distinct congenic background to allow for identification of donor populations. For experiments with *in vivo* T_ex_, mice were sacrificed between 22-31dpi with LCMV-Cl13.

### Infections

LCMV strains Armstrong (Arm) and clone 13 (Cl13) were propagated and titers were determined as previously described (*6*). C57BL/6 mice were infected intraperitoneally (i.p.) with 2×10^5^ plaque-forming units (PFU) of LCMV-Arm or i.v. with 4×10^6^ PFU LCMV-Cl13.

### *In vitro* chronic and acute stimulation

CD8 T cells were isolated from spleens of naïve P14 transgenic mice by homogenizing against a 70μm cell strainer. The cell suspension was then washed and passed through a 70μm cell strainer an additional time. CD8 T cells were isolated via negative selection kit (StemCell) per manufacturer’s instructions. Dendritic cells (DCs) were isolated from spleens of naïve WT C57BL/6 mice by cutting samples into small pieces and incubating at 37°C for 45 minutes with 1U/ml DNase I (Roche) and 10U/ml collagenase D (Roche) in complete RPMI (cRPMI): RPMI 1640 (Corning Cellgro) supplemented with 10% FBS, 50μM β-mercaptoethanol, 2mM glutamine (Gibco), 100U/ml penicillin (Sigma), 100μg/ml streptomycin (Sigma), 100μM non-essential amino acids (Invitrogen), 1mM sodium pyruvate (Invitrogen), and 20mM HEPES buffer (Invitrogen). Cells were then homogenized against a 70μm cell strainer and red blood cells were lysed in ACK lysis buffer (Gibco) for 5 minutes. The cell suspension was quenched and washed with PBS then passed through a 70μm cell strainer an additional time. DCs were isolated with CD11c+ Microbeads (Miltenyi) per manufacturer’s instructions then pulsed with D^b^GP^33-41^ peptide (0.1μM, GenScript) in cRPMI for 30 minutes at 37°C. CD8 P14 cells and DCs were then co-cultured in cRPMI at a 1:1 ratio (10^5^ DCs + 10^5^ P14 cells per well) in a flat-bottom 96-well plate as previously described (*15*). At days 2, 4, and 6 of co-culture, P14 cells were counted via BD Accuri, normalized to a concentration of 0.5×10^6^ P14 cells/ml (10^5^ P14 cells per well), and re-plated into *in vitro* culture. Chronically stimulated P14 cells received D^b^GP^33-41^ peptide (1nM) at days 2, 4, and 6; acutely stimulated P14 cells received initial peptide stimulation but no subsequent doses of peptide at days 2, 4, and 6. Both *in vitro* conditions received 10U/ml recombinant human IL-2 (Peprotech) at days 2, 4, and 6.

*In vitro* chronic and acute stimulation via αCD3/αCD28 was performed as previously described (*18*).

### Flow cytometry

*Ex vivo* single cell suspensions were prepared by homogenizing against a 70μm cell strainer followed by red blood cell lysis with ACK lysis buffer (Gibco). *In vitro* single cell suspensions were prepared by resuspending cells in well with multichannel pipette. Cells were washed with staining buffer (PBS with 2% FCS) and surface staining was performed for 1 hour at 4°C in staining buffer. For intranuclear staining, permeabilization was performed using the Foxp3/Transcription Factor Fixation/Permeabilization kit (eBioscience) for 30 minutes at 4°C, following which TF staining was performed for 2 hours at 4°C. For intracellular cytokine staining (ICS), permeabilization was performed using the BD Cytofix/Cytoperm™ Fixation/Permeabilization Solution Kit (BD) for 30 minutes at 4°C, following which ICS was performed for 1 hour at 4°C. Samples were run on a BD LSR II. Voltages on the machine were standardized using fluorescent targets and rainbow beads (Spherotech). Data were analyzed with FlowJo software (v10.7.1, TreeStar). Cell sorting was performed on a BD FACSAria with a 100μm nozzle; a circulating temperature system was used at 4°C when sorting for sequencing and 37°C when sorting for *in vivo* transfer. A small aliquot of all sorted samples was run as a purity check.

### Preparation of RNA- and ATAC-seq libraries

At day 7 of *in vitro* culture, live CD8 T cells were sorted to a purity of >95% for each replicate. Naïve CD8 T cells were also sorted from P14 mice. To extract RNA, 5×10^4^ cells were resuspended in buffer RLT supplemented with β-mercaptoethanol and processed with an RNeasy Micro Kit (Qiagen) per manufacturer’s instructions. mRNA libraries were prepared using the SMART-Seq v4 Ultra Low Input RNA Kit (Takara) per manufacturer’s instructions. Extracted RNA and libraries were assessed for quality on a TapeStation 2200 instrument (Agilent). ATAC libraries were generated as described with minor changes (*85*). Briefly, nuclei from 5×10^5^ cells were isolated using a lysis solution composed of 10mM Tris-HCl, 10mM NaCl, 3mM MgCl2, and 0.1% IGEPAL CA-630. Immediately following cell lysis, nuclei were pelleted in DNA Lo-Bind 1.5ml tubes (Eppendorf) and resuspended in TD Buffer with Tn5 transposase (Illumina). Transposition reaction was performed at 37°C for 45 minutes. DNA fragments were purified from enzyme solution using MinElute Enzyme Reaction Cleanup Kit (Qiagen). Libraries were barcoded (Nextera Index Kit, Illumina) and amplified with NEBNext High Fidelity PCR Mix (New England Biolabs). Library quality was assessed using a TapeStation instrument. RNA and ATAC libraries were quantified using a KAPA Library Quantification Kit and sequenced on an Illumina NextSeq 550 instrument (150bp, paired-end) on high-output flow cells.

### RNA-seq data processing and analysis

FASTQ files were aligned using STAR (v2.5.2a) against the mm10 mouse reference genome. The aligned files were processed using PORT gene-based normalization (https://github.com/itmat/Normalization). Computational merging of *in vitro* and *in vivo* data sets was performed using the Combat function in the sva R package (v3.40.0). Differentially expressed genes (DEGs) were identified with DESeq2 (v1.32.0, DESeq function) using log_2_ fold change > 1 and adjusted p value < 0.05; genes were first filtered on minimum expression (median 20 reads per group). GSEA (*39, 40*) and GSVA (*45*) were used to calculate enrichment scores; an FDR of 0.25 was used to determine statistical significance in GSEA.

### ATAC-seq data processing and analysis

The script used for processing raw ATAC-seq FASTQ data is available at the following GitHub repository: https://github.com/wherrylab/jogiles_ATAC/blob/master/Giles_Wherry_ATAC_pipeline_mm10_UPennCluster. In brief, samples were aligned to mm10 reference genome with Bowtie2 (v2.1.0). Unmapped, unpaired, and mitochondrial reads were removed using samtools (v1.1). ENCODE Blacklist regions were removed (https://sites.google.com/site/anshulkundaje/projects/blacklists). PCR duplicates were removed using Picard (v1.141). Peak calling was performed with MACS2 (v2.1.1) with an FDR q-value = 0.01. A union peak list for each experiment was created by combining all peaks in all samples; peaks had to be present in 2 of 4 technical replicates in order to be called. Overlapping peaks were merged using bedtools (v2.29.2) merge. The number of reads in each peak was determined with bedtools (v2.29.2) coverage. Peaks were annotated using HOMER (v4.11.1). Computational merging of *in vitro* and *in vivo* data sets was performed using the Combat function in the sva R package (v3.40.0). Differentially accessible chromatin regions (DACRs) were identified with the R package DESeq2 (v1.32.0, DESeq function) using log_2_ fold change > 1 and adjusted p value <0.05. ATAC-seq signal tracks were generated with the gviz R package (v1.36.1). ATAC signal tracks are generated using gviz with bigwigs normalized for library size and group scaled across all samples within each dataset; peaks out of range are indicated with a contrast color on top. Scripts for peak set enrichment are available at: https://github.com/wherrylab/jogiles_ATAC/blob/master/PSEA.Rmd. In brief, bedtools (v2.29.2) intersect was used to find overlapping peaks between the experiment and peak set of interest. Peak names between the experiment and peak set of interest were unified using custom R scripts. GSEA (*39, 40*) was used to calculate enrichment scores; an FDR of 0.25 was used to determine statistical significance.

### Transcriptional and epigenetic trajectory analysis

To evaluate transcriptional and epigenetic trajectories, genes were manually binned into patterns based on changes in gene expression and chromatin accessibility between days 0 and 4, and days 4 and 7 of *in vitro* culture. A log_2_ fold change > 1 between two timepoints was scored as 1; a log_2_ fold change < −1 was scored as −1; a log_2_ fold change between 1 and −1 was scored as 0. The relative gene expression and chromatin accessibility was determined by starting all genes at a baseline of 0, then adding these scores over time.

### Taiji analysis

Taiji (v1.3) analysis was implemented as previously described (http://wanglab.ucsd.edu/star/taiji) (*52, 53*). Normalized ranks were compared across conditions and top TFs were chosen using cut-offs of mean > 0.0001 and fold change of 5 above the mean. TFs were visualized using the pheatmap R package (v1.0.12).

### TF co-occurrence analysis

The TF-COMB python package (https://tf-comb.readthedocs.io/en/latest/) (*86*) was used to find TF binding sites in ACRs using the CIS-BP motif library of TFs (cisbp.ccbr.utoronto.ca) for mm10. Market basket analysis was done on these TFBSs to identify and count co-occurring transcription factors using the default window of 100bp and no allowed overlap. TFs were then filtered on minimum gene expression from RNAseq (median 20 reads per group).

### Pooled CRISPR screening

#### Vector and library construction

SpCas9 sgRNA was expressed using pSL21-VEX (U6-sgRNA-EFS-VEX, available through Addgene). 4-5 sgRNA were designed against functional domains of each TF (*56*), synthesized by Integrated DNA Technologies (IDT), pooled in equal molarity, PCR-amplified, and cloned into BsmBI-digested SL21 vector as previously described (*57*).

#### In vitro retroviral transduction

CD8 T cells were isolated from spleens of naïve C9P14 transgenic mice as described above. Total splenocytes from naïve WT C57BL/6J mice were isolated via collagenase/DNase digestion as described above. Total splenocytes were pulsed with D^b^GP^33-41^ peptide (0.1μM, GenScript) in cRPMI for 30 minutes at 37°C and then co-cultured with C9P14 cells at a 10:1 ratio (10^6^ splenocytes + 10^5^ C9P14 cells per well) in a flat-bottom 96-well plate. 24 hours after initial stimulation, C9P14 cells were enriched using an EasySep magnetic CD8 negative selection kit (StemCell) per manufacturer’s instructions and spin-transduced for 60 minutes at 2000xg 37°C with RV supernatant containing polybrene (4mg/mL) and IL-2 (100U/mL) (*75*). 6 hours later, C9P14 cells were returned to *in vitro* co-culture with either D^b^GP^33-41^ pulsed or unpulsed splenocytes for *in vitro* chronic and acute stimulation conditions, respectively. Chronic and acute stimulation *in vitro* proceeded as described above, with cells being counted, normalized to a concentration of 0.5×10^6^ C9P14 cells/ml (10^5^ C9P14 cells per well), and put back into culture either with or without D^b^GP^33-41^ peptide at days 3 and 5. A portion of transduced C9P14 cells were kept in *in vitro* culture without splenocyte co-culture; 24 hours after transduction, sgRNA^+^Cas9^+^ cells were sorted from this population for baseline sequencing. sgRNA^+^Cas9^+^ cells were sorted from *in vitro* chronically and acutely stimulated P14 populations for sequencing at day 7.

#### Isolated library construction, sequencing, and data processing

Library preparation proceeded as previously described (*57*). Read counts of each individual sgRNA were calculated and compared to the sequence of reference sgRNA as previously described (*56*). Significant hits were calculated via MAGeCK (v0.5.8) (*87*) with p < 0.05 and log_2_ fold change < 0.5.

#### GSEA comparing in vitro and in vivo CRISPR screening

Gene signatures were created from significantly enriched and depleted hits (significance calculated via MAGeCK (v0.5.8) (*87*) with p < 0.1) from previously published *in vivo* CRISPR screening (*57*) in spleen at 14dpi with LCMV-Cl13. GSEA (*39, 40*) was used to calculate enrichment of these gene signatures in the total ranked order gene set from CRISPR screening in *in vitro* chronically stimulated P14 cells.

### shRNA and overexpression vector generation

shRNA vector against *Bhlhe40* was generated as previously described (*88, 89*) using guide sequence TAGGATACACTTGAAATCCGTG. Overexpression plasmid encoding *Bhlhe40* was kindly gifted by M. M. Gubin (University of Texas MD Anderson Cancer Center). RV was generated for each construct as previously described (*75*) using 293T cells (ATCC).

### *In vivo* retroviral transduction

Single cell suspensions were prepared by mechanically disrupting spleens from congenically distinct P14 mice by homogenizing against a 70μm cell strainer. CD8 T cells were enriched using an EasySep magnetic negative selection kit (StemCell) per manufacturer’s instructions. P14 cells were stimulated with αCD3 (1mg/mL; Invitrogen), αCD28 (0.5mg/mL; Invitrogen), and recombinant human IL-2 (100U/ml; Peprotech) in cRPMI. 24 hours post activation, P14 cells were spin-transduced for 60 minutes at 2000xg 37°C with RV supernatant containing polybrene (4mg/mL) and IL-2 (100U/mL) (*75*). Approximately 24 hours later, GFP+ cells were sorted on a BD FACSAria II. Sorted cells were washed with and resuspended in warmed PBS. P14 cells transduced with control (*Krt8* shRNA or empty OE vector) or experimental RV (*Bhlhe40* shRNA or *Bhlhe40* OE vector) were pooled at a 1:1 ratio (2.5×10^4^ control + 2.5×10^4^ *Bhlhe40* KD/OE P14 cells per mouse) and transferred i.v. into congenically distinct recipient mice infected with LCMV-Arm or -Cl13 one day prior.

### αPD-L1 treatment

αPD-L1 (clone 10F.9G2; BioXcell) treatment (200ug/mouse) was administered i.p. every 3 days from 22dpi-35dpi for a total of 5 treatments (*2*); PBS was used as a vehicle control.

### Immunoblots

2×10^6^ cells per condition were sorted. Whole-cell lysates were prepared in RIPA buffer and separated on a 4-15% gel by SDS-PAGE, transferred to a PVDF membrane and blocked in 5% non-fat milk in TBST. Primary antibodies against the following proteins were used: GAPDH (1:1000; CST) and Bhlhe40 (1:500; Novus Biologicals) in 1% non-fat milk in TBST. HRP-conjugated anti-rabbit secondary antibodies were used (CST). Signal was detected using SuperSignal West Femto ECL substrate (Thermo) and imaged on a BioRad ChemiDoc Touch imaging system.

### Statistical analyses

GraphPad Prism software was used for all statistical analyses. Statistics were calculated by paired two-tailed t test. The statistical details for each experiment are provided in the associated figure legends.

## Acknowledgements

We thank all members of the Wherry laboratory for helpful discussions and critical analysis of the manuscript. Figure schematics were created with Biorender.com.

## Author Contributions

Conceptualization: JEW, EJW

Methodology: JEW, ZC, JS, EJW

Resources: SM, JRG, SFN, AEB, ZC, JS

Investigation: JEW, JRG, MLC, EF, JHL

Formal Analysis: JEW, SM, JRG, HH, JS

Visualization: JEW, SM, JRG, HH

Writing – original draft: JEW, EJW

Writing – review & editing: JEW, JRG, SFN, AEB, ZC, OK, RPS, YJH

Funding Acquisition: EJW, JS

Supervision: EJW, JS

## Code and data availability

RNA- and ATAC-seq data have been deposited in the NCBI Gene Expression Omnibus (GEO) database and are accessible through the GEO SuperSeries accession number GSE211015. All other relevant data are available from the corresponding author upon reasonable request.

---

**Fig. S1.**
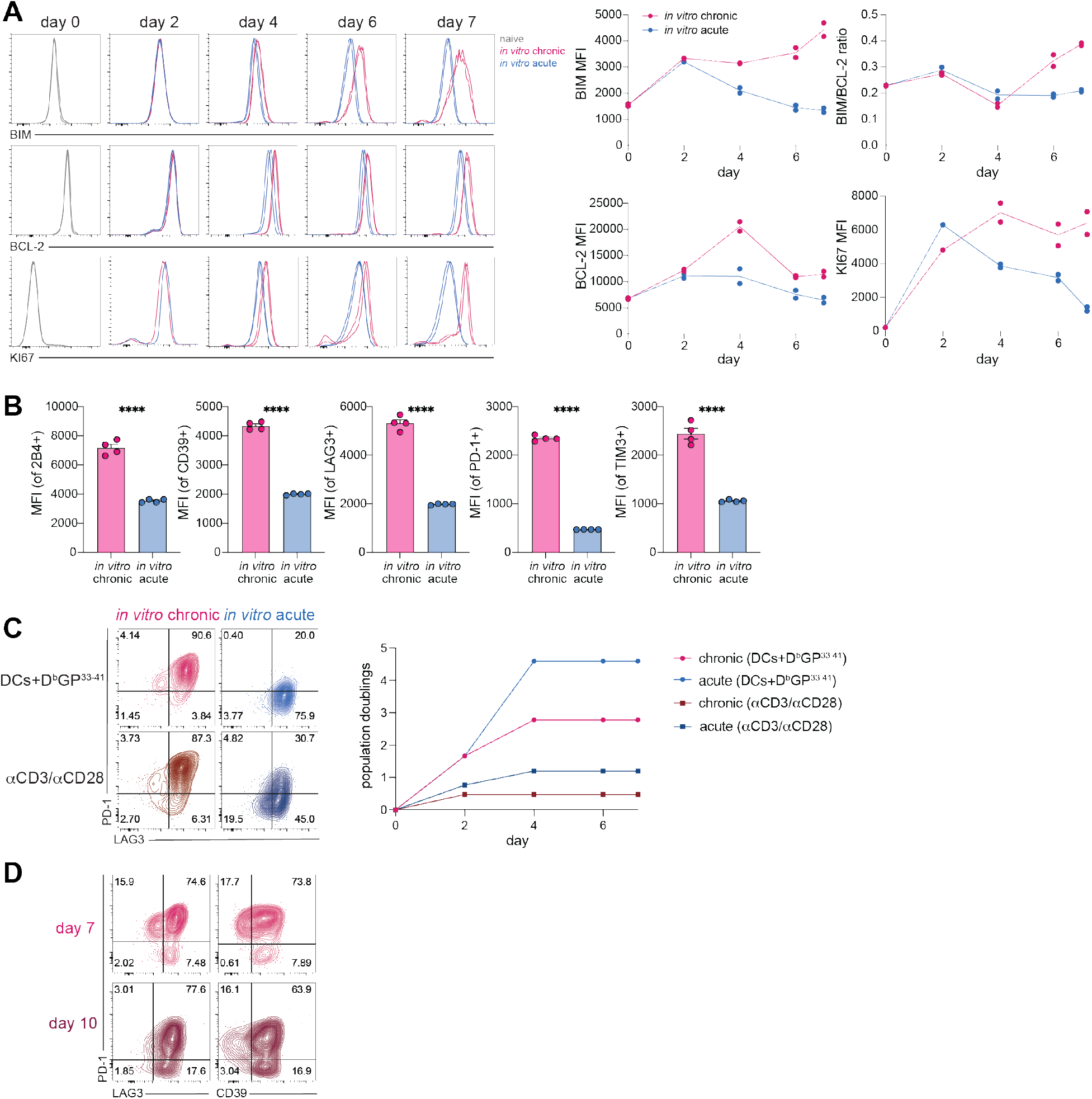
Chronic antigenic stimulation *in vitro* induces high per-cell IR expression. **(A)** [left] Histograms and [right] summary data of longitudinal expression of BIM, BCL-2, and KI67 by *in vitro* chronically and acutely stimulated P14 cells. **(B)** Summary data indicating MFI (gated on IR+ population of CD44^hi^ CD8+ live singlets) of IRs after chronic and acute stimulation of P14 cells *in vitro*. Significance calculated by unpaired two-tailed t test; **\*\*\*\***p< 0.0001. **(C)** [left] Representative flow cytometry data of IR expression (gated on CD44^hi^ CD8+ live singlets) and [right] cell expansion after chronic and acute stimulation, either via DCs+D^b^GP^33-41^ or αCD3/αCD28. **(D)** Representative flow cytometry data of IR expression (gated on CD44^hi^ CD8+ live singlets) after chronic stimulation for either 7 or 10 days.

**Fig. S2.**
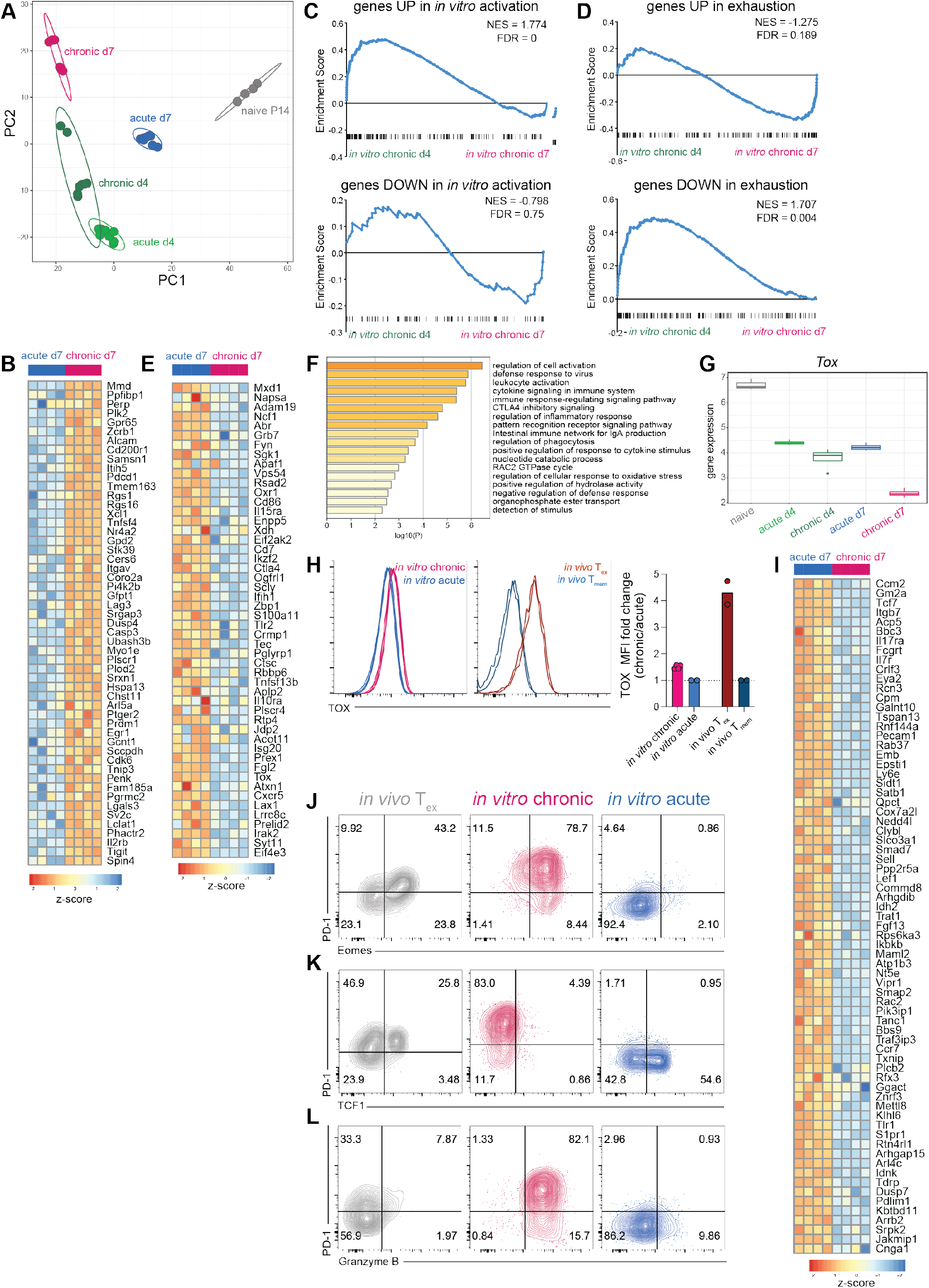
*In vitro* chronically stimulated P14 cells develop a transcriptional signature of T_ex_. **(A)** PCA of RNA-seq data from *in vitro* chronically and acutely stimulated P14 cells. **(B)** Gene expression of leading edge genes upregulated in *in vivo* T_ex_ enriched after chronic stimulation *in vitro*. **(C-D)** GSEA of gene sets for **(C)** *in vitro* activation and **(D)** exhaustion (*41*) in *in vitro* chronically stimulated cells at d4 and d7. **(E)** Heatmap and **(F)** GO analysis of genes upregulated in *in vivo* T_ex_ that were not enriched after chronic stimulation *in vitro*. **(G)** Gene expression of *Tox* in *in vitro* chronically and acutely stimulated P14 cells. **(H)** [left] TOX protein expression in *in vitro* chronically and acutely stimulated P14 cells and *in vivo* T_ex_ and T_mem_. [right] Fold change of TOX MFI in *in vitro* chronic over acute and *in vivo* T_ex_ over T_mem_. **(I)** Gene expression of leading edge of genes downregulated in *in vivo* T_ex_ enriched after chronic stimulation *in vitro*. **(J-L)** Representative flow cytometry data detailing co-expression of PD-1 and **(J)** Eomes, **(K)** TCF1, and **(L)** Granzyme B on *in vivo* T_ex_ (LCMV-Cl13 30dpi), *in vitro* chronically stimulated P14 cells, and *in vitro* acutely stimulated P14 cells (all gated on CD44^hi^ CD8+ live singlets). Numbers in flow cytometry plots indicate percentage of parent population within each gate.

**Fig. S3.**
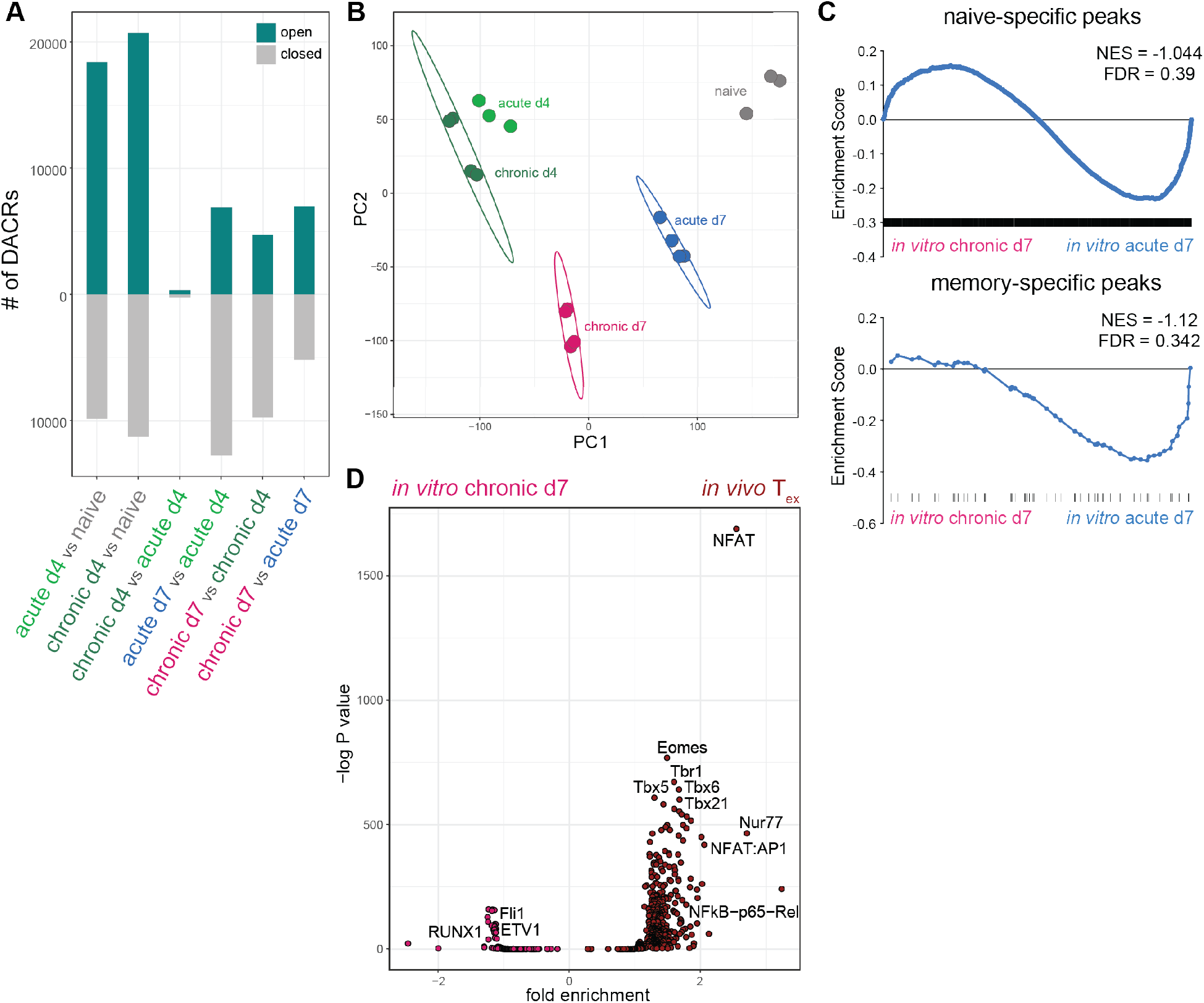
*In vitro* chronically stimulated P14 cells develop epigenetic signatures of T_eff_ and T_ex_. **(A)** Number of DACRs (filtered on lfc>1 and p<0.05) between pairwise comparisons as indicated. **(B)** PCA of ATAC-seq data from *in vitro* chronically and acutely stimulated P14 cells. **(C)** PSEA of naive- and memory-specific ACRs in *in vitro* chronically and acutely stimulated P14 cells. **(D)** TF motif accessibility in DACRs between *in vitro* chronically stimulated P14 cells and *in vivo* T_ex_.

**Fig. S4.**
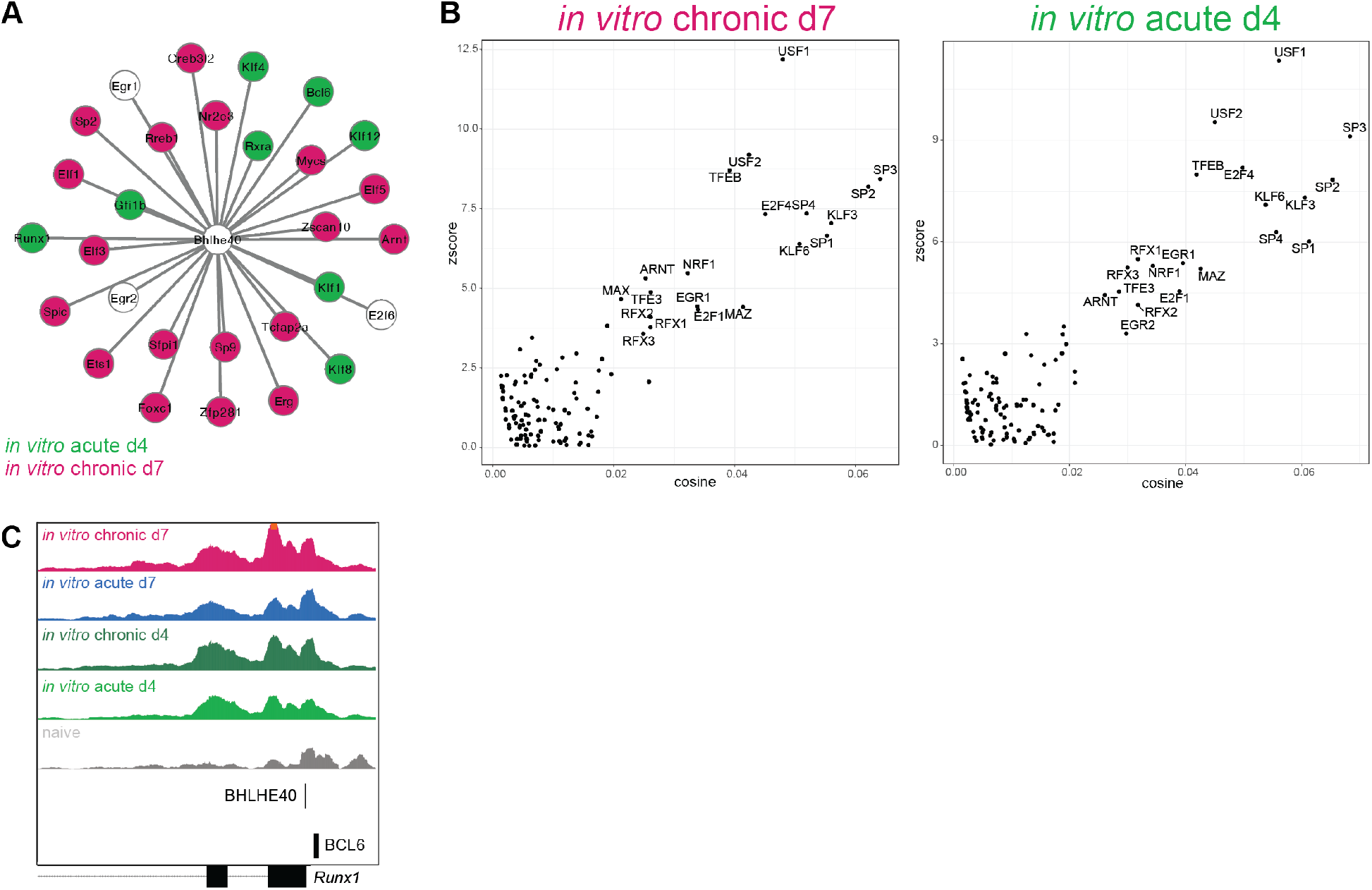
BHLHE40 is predicted to regulate known transcriptional networks associated with CD8 T cell exhaustion. **(A)** Taiji analysis of differential transcriptional networks downstream of BHLHE40 at d7 of chronic stimulation *in vitro* and d4 of acute stimulation *in vitro*. Shared downstream TFs indicated in white. **(B)** TF motifs that co-occur with BHLHE40 motifs. **(C)** ATAC-seq signal tracks for *in vitro* chronically and acutely stimulated P14 cells for the *Runx1* locus; binding motifs for BHLHE40 and BCL6 shown below.

**Fig. S5.**
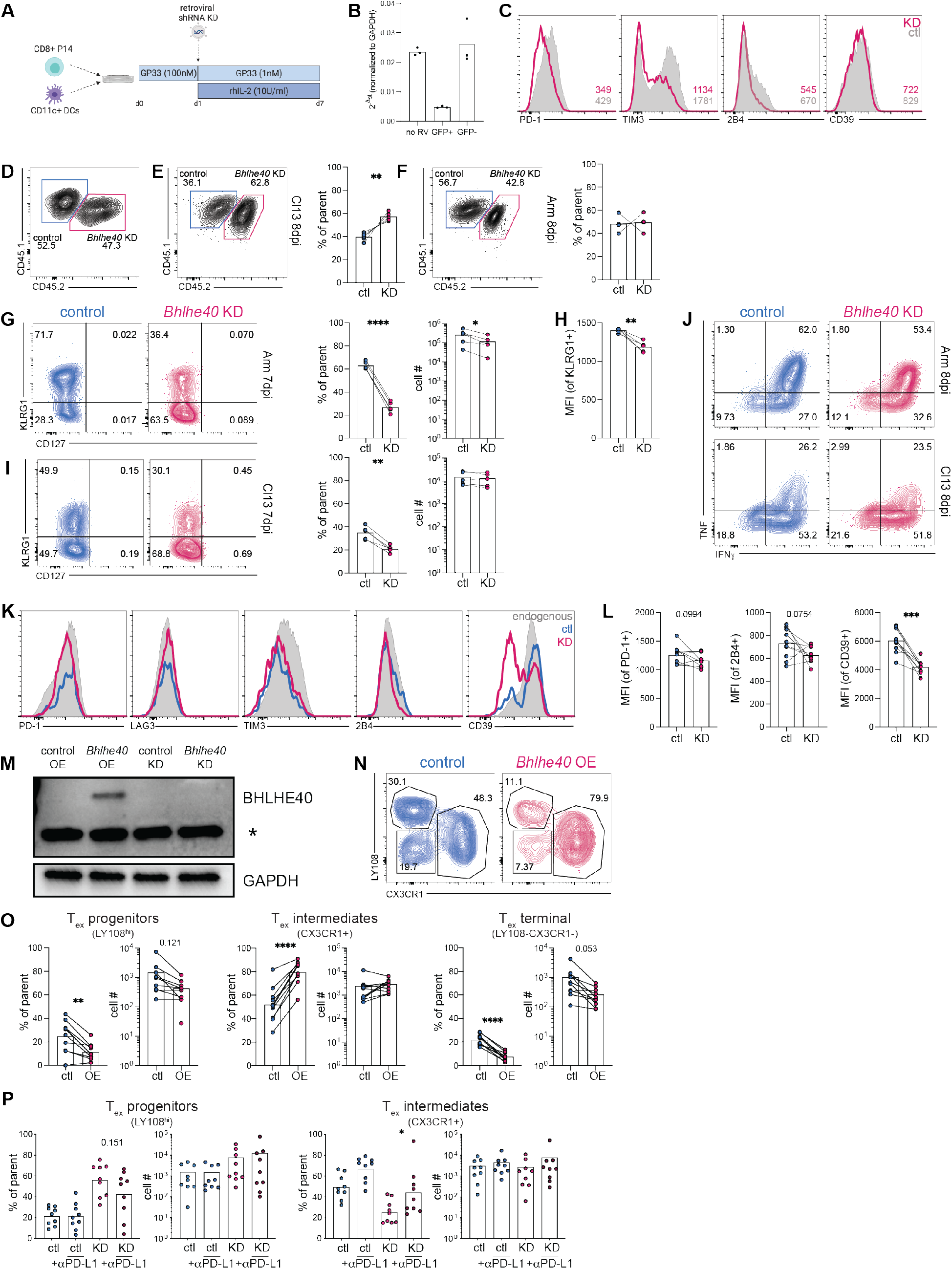
BHLHE40 is a novel transcriptional regulator of CD8 T cell differentiation. **(A)** Experiment schematic of shRNA KD of *Bhlhe40* in *in vitro* chronically stimulated P14 cells. **(B)** Relative mRNA expression after *in vitro Bhlhe40* KD. **(C)** Representative flow cytometry histograms detailing IR expression by control and *Bhlhe40* KD P14 cells (gated on GFP+ CD44^hi^ CD8+ live singlets) after chronic stimulation *in vitro*. Mean fluorescence intensity (MFI) of each population indicated in lower right; representative of 2 experiments. **(D)** Representative flow cytometry plot of control and *Bhlhe40* KD input P14 cells (gated on GFP+ CD44^hi^ CD8+ live singlets) before transfer. **(E,F)** [left] Concatenated flow cytometry plot and [right] summary data of frequency of control and *Bhlhe40* KD P14 cells at 8dpi of **(E)** LCMV-Cl13 and **(F)** LCMV-Arm. **(G,I)** [left] Concatenated flow cytometry plots and [right] summary data of frequency and total number of KLRG1+ control and *Bhlhe40* KD P14 cells at 7dpi of **(G)** LCMV-Arm or **(I)** LCMV-Cl13. **(H)** MFI of KLRG1 (gated on KLRG1+ population) in control and *Bhlhe40* KD P14 cells at 7dpi of LCMV-Arm. **(J)** Concatenated flow cytometry plots of effector cytokine production in control and *Bhlhe40* KD P14 cells at 8dpi of [top] LCMV-Arm and [bottom] -Cl13. **(K)** Concatenated flow cytometry histograms detailing IR expression on endogenous (gated on GFP-gp33+ CD44^hi^ CD8+ live singlets), control, and *Bhlhe40* KD P14 cells at 31dpi of LCMV-Cl13. **(L)** Summary data indicating MFI (gated on IR+ population) of IRs on control and *Bhlhe40* KD P14 cells at 31dpi of LCMV-Cl13. **(M)** Protein expression of BHLHE40 after *in vitro* overexpression (OE) and KD of *Bhlhe40*; * indicates non-specific binding. **(N)** Concatenated flow cytometry plots and **(O)** summary data of frequency of T_ex_ subsets in control and *Bhlhe40* OE P14 cells at 31dpi of LCMV-Cl13. **(P)** Frequency and total number of progenitor and intermediate T_ex_ in control and *Bhlhe40* KD P14 cells at 37dpi of LCMV-Cl13, after treatment with vehicle control or αPD-L1 from 22-35dpi. **(E-L,N-P)** n=5 mice (Arm), n=10 mice (Cl13), representative of 3 experiments. Significance calculated by paired two-tailed t test; **\***p<0.05, **\*\***p<0.01, **\*\*\***p<0.001, **\*\*\*\***p<0.0001. **(D-J,N)** Numbers in flow cytometry plots indicate percentage of parent population within each gate.

